# CBL mutations promote activation of PI3K/AKT signaling via LYN kinase

**DOI:** 10.1101/2020.04.13.038448

**Authors:** Roger Belizaire, Sebastian H.J. Koochaki, Namrata D. Udeshi, Alexis Vedder, Lei Sun, Tanya Svinkina, Christina Hartigan, Caroline Stanclift, Monica Schenone, Steven A. Carr, Eric Padron, Benjamin L. Ebert

## Abstract

*CBL* encodes an E3 ubiquitin ligase and signaling adaptor that acts downstream of cytokine receptors. Recurrent *CBL* mutations occur in myeloid malignancies, but the mechanism by which these mutations drive oncogenesis remains incompletely understood. Here we performed a series of studies to define the phosphoproteome, CBL interactome and molecular mechanisms of signaling activation in cells expressing an allelic series of *CBL* mutants. Our analyses revealed that increased LYN activation and interaction with mutant CBL are key drivers of enhanced PIK3R1 recruitment and downstream PI3K/AKT signaling in *CBL*-mutant cells. Furthermore, we demonstrated *in vitro* and *in vivo* efficacy of LYN inhibition by dasatinib in *CBL*-mutant cell lines and primary chronic myelomonocytic leukemia cells. Overall, our data provide rationale for exploring the therapeutic potential of LYN inhibition in patients with *CBL*-mutated myeloid malignancies.

**Statement of Significance:** We investigated the oncogenic mechanisms of myeloid malignancy-associated *CBL* mutations by mass spectrometry-based proteomics and interactomics. Our findings indicate that increased LYN kinase activity in *CBL*-mutant cells stimulates PI3K/AKT signaling, revealing opportunities for the use of targeted inhibitors in *CBL*-mutated myeloid malignancies.

## Introduction

Hematologic malignancies are often characterized by somatic alterations in genes encoding signaling proteins, leading to increased cytokine sensitivity, cell survival and proliferation (1). Recurrent somatic mutations in the *<u>C</u>asitas <u>B</u>-lineage <u>L</u>ymphoma* (*CBL*) gene, which encodes an E3 ubiquitin ligase and signaling adaptor, occur in myelodysplastic syndromes (MDS) and other myeloid neoplasms (2–9), including 10-20% of chronic myelomonocytic leukemia (CMML) patients (10, 11). In addition, up to 20% of children diagnosed with juvenile myelomonocytic leukemia (JMML) harbor germline *CBL* mutations (12–15). The presence of *CBL* mutations has been associated with poor prognosis (16–18), and a better understanding of the mechanisms by which *CBL* mutations promote myeloid disease is needed for the development of new and effective therapeutic strategies.

The domain structure of the CBL protein comprises an N-terminal tyrosine kinase binding (TKB) domain followed by a linker region, RING domain, proline-rich region (PRR), C-terminal phosphorylation sites, and ubiquitin association domain (19). As a signaling adaptor, CBL’s TKB domain, PRR and C-terminal phosphorylation sites facilitate coupling between tyrosine-phosphorylated cytokine receptors on the cell membrane and intracellular proteins involved in signal transduction. The RING domain, which binds E2 ubiquitin ligase proteins, and linker region are both essential for CBL’s E3 ubiquitin ligase function. The majority of *CBL* mutations in myeloid malignancies are predicted to alter the ubiquitin ligase activity of CBL through single amino acid substitutions within the linker region or RING domain (3,5–15), or splice site alterations resulting in exclusion of most amino acids within the RING domain (15, 20). Recurrent mutations predicted to affect CBL’s signaling adaptor functions are exceptionally rare, implying that oncogenic *CBL* mutations result in selective loss of CBL’s E3 ubiquitin ligase function. Moreover, *CBL* mutations often occur in the setting of acquired 11q uniparental disomy including the *CBL* locus, suggesting that expression of wild-type (WT) *CBL* impairs the oncogenic phenotype in cells expressing mutant *CBL* (3,5,6,21). Consistent with the genetics of *CBL* mutations in myeloid malignancies, missense mutations in the RING domain of murine *Cbl* promoted the development of a myeloproliferative disorder that was not observed in *Cbl* knockout mice (22–24). Together with work from other groups (6, 15), these data indicate that *CBL* mutations confer a gain-of-function associated with disruption of E3 ubiquitin ligase activity and preservation of signaling adaptor functions (21).

Since *CBL* mutations inactivate the ubiquitin ligase activity of CBL while retaining domains needed for downstream signaling, we hypothesized that defective degradation of phosphorylated proteins causes oncogenic activation of signaling pathways. To identify these pathways, we performed unbiased and comprehensive analyses of the signaling pathways and CBL interactome in a panel of cell lines expressing an allelic series of *CBL* mutations using quantitative liquid chromatography-tandem mass spectrometry (LC-MS/MS). We found that the CBL-LYN-PI3K axis (25–28) plays a significant role in the gain-of-function phenotype conferred by *CBL* mutations. In addition, LYN inhibition by dasatinib effectively diminished the expansion of *CBL* mutant cell lines and primary CMML samples *in vitro* and *in vivo*, highlighting the potential of LYN-targeted therapies in patients with *CBL*-mutated myeloid disease.

## Results

### Characteristics of *CBL* mutations in 191 patients

To select common, recurrent *CBL* mutations for functional studies, we examined clinical sequencing data from 9122 patients undergoing evaluation for hematologic disorders, including MDS and myeloid leukemias (29). Two-hundred fourteen *CBL* mutations were detected (Supplementary Figure 1A), of which one-hundred ninety-five (91%) were single nucleotide changes leading to amino acid substitutions predicted to disrupt the function of the linker region or RING domain (Supplementary Figure 1B). Overall, single amino acid substitutions at positions Y371, L380, C384, C404 and R420 were most common, comprising nearly 50% of all *CBL* mutations (Supplementary Figure 1C).

### CBL mutations activate LYN and PI3K/AKT signaling pathways

To elucidate the mechanisms by which mutant *CBL* promotes myeloid oncogenesis, we generated an allelic series of *CBL* mutations that reflects the genetics of human myeloid malignancies in cytokine-dependent cell lines. We focused on our analysis on three common *CBL* missense mutations, one that occurs within the linker region (Y371H) and two that occur within the RING domain (C384Y and R420Q). To generate an *in vitro* model consistent with the observed 11q UPD observed in patients with *CBL* mutations (3,5,6,21), we first employed CRISPR-Cas9 to produce *Cbl* knockout clones in the murine myeloblast cell line, 32D (Supplementary Figure 2), which proliferates in response to either IL3 or GM-CSF. To investigate the effects of *CBL* mutations on cytokine signaling and proliferation, we next generated a panel of cell lines expressing human, V5-tagged, mutant (i.e., Y371H, C384Y or R420Q) or WT *CBL* in 32D-*Cbl*^KO^ cells. Compared to expression of *CBL* WT, expression of *CBL* mutants was associated with increased IL3 and GM-CSF sensitivity in cell proliferation assays (Figures 1A and B); expression of the same *CBL* mutants in the murine lymphoblast cell line BaF3 also led to increased IL3 sensitivity and was sufficient to promote IL3-independence (Figures 1C and Supplementary Figure 3).

**Figure 1.**
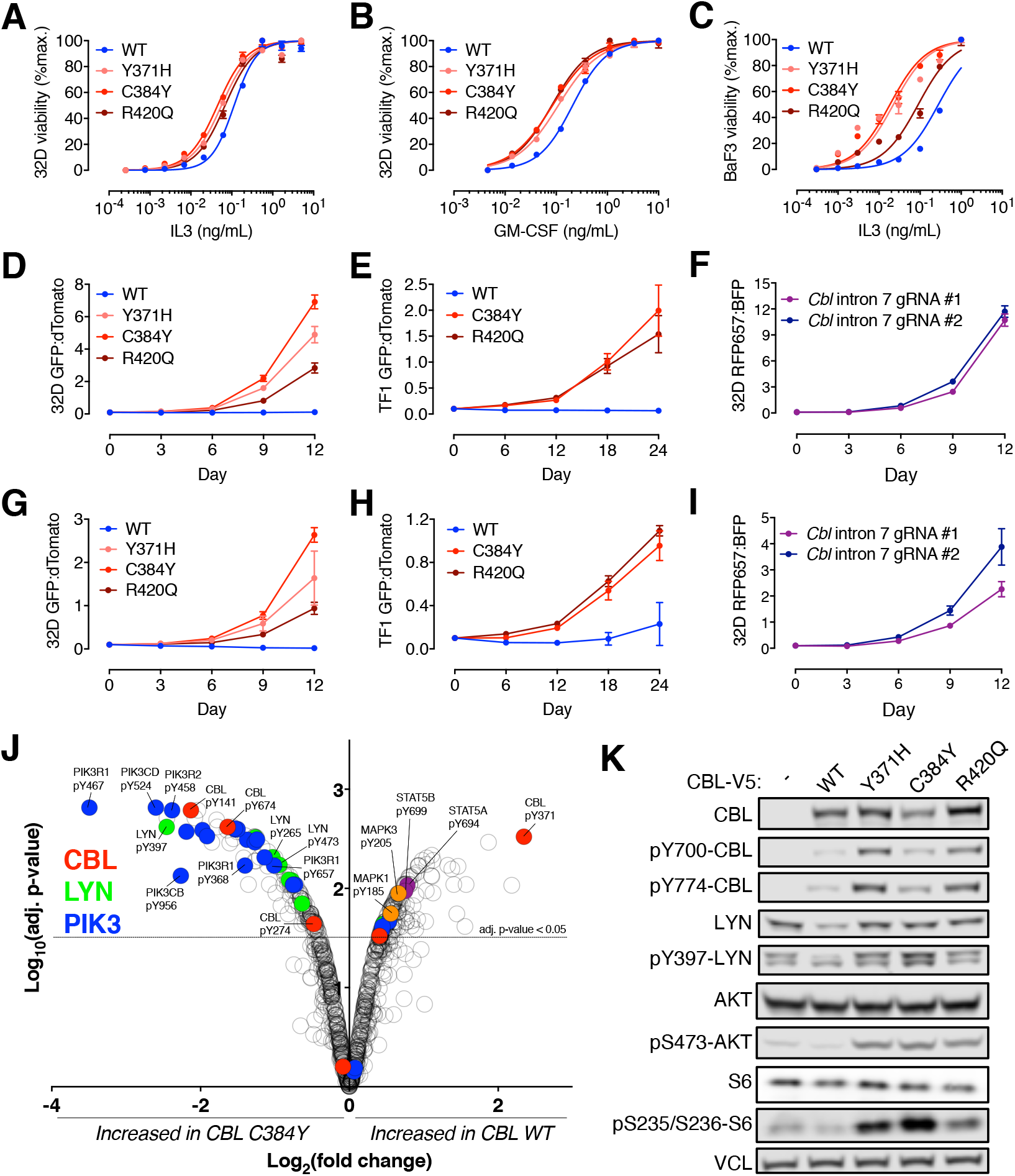
Gain-of-function *CBL* mutations are associated with increased cytokine sensitivity and activation of LYN and PI3K/AKT signaling pathway. (A-C) Proliferation of *Cbl*^KO^ 32D (A and B) or BaF3 (C) cells expressing WT or mutant CBL after 3-day stimulation with IL3 (A and C) or GM-CSF (B). (D-I) Competition between GFP-labeled 32D-*Cbl*^KO^ (D and G) or TF1-*CBL*^KO^ (E and H) cells expressing CBL mutant and dTomato-labeled CBL WT (D and E) or CBL KO (G and H) cells. Competition between 32D-Cas9 cells expressing *Cbl* intron 7 gRNAs (RFP657) and non-targeting gRNA (BFP, panel F) or *Cbl* exon 1 gRNAs (BFP, panel I). (J) Volcano plot showing relative quantity of phosphotyrosine sites detected by mass spectrometry and normalized to proteome level data in 32D-*Cbl*^KO^ cells expressing V5-tagged CBL WT or C384Y. (K) Western blot for total and phosphorylated CBL, LYN, AKT, and S6 proteins in 32D-*Cbl*^KO^ cells expressing V5-tagged CBL WT, Y371H, C384Y, or R420Q. Western blot for vinculin (VCL) was used as a loading control.

To determine whether cells expressing *CBL* mutants had a competitive advantage over cells expressing *CBL* WT, we co-cultured GFP-labeled *CBL* mutant cells with dTomato-labeled *CBL* WT cells at a ratio of 1:10 in limiting IL3 (0.1ng/mL) and performed serial measurements of the GFP:dTomato ratio using flow cytometry. For each *CBL* mutant tested, the GFP:dTomato ratio increased significantly over several days, indicating that expression of *CBL* mutants conferred a proliferative advantage (Figure 1D); similar results were observed in the GM-CSF-dependent, human cell line, TF1 and in BaF3 (Figure 1E and Supplementary Figures 3B). Cells expressing *CBL* mutants also displayed a proliferative advantage over cells with complete loss of *CBL* (i.e., 32D-*Cbl*^KO^ or TF1-*CBL*^KO^), consistent with the expected gain-of-function of the CBL mutant protein (Figures 1G and H). Finally, CRISPR-Cas9-mediated editing of endogenous *Cbl* exon 8 splice sites in 32D cells led to exon 8 exclusion, similar to *CBL* exon 8 splice site mutations in human myeloid disease (15, 20), as well as increased cell proliferation compared to both *Cbl*^WT^ and *Cbl*^KO^ 32D cells (Supplementary Figure 4 and Figures 1F and I). Altogether, these mouse and human cell lines model myeloid disease-associated *CBL* mutations by demonstrating that selective disruption of CBL’s linker region or RING domain leads to gain-of-function effects on cytokine sensitivity and cell proliferation.

We leveraged our *in vitro* models of *CBL* mutations to explore the signaling pathways that are activated in *CBL* mutant cells. Given the role of CBL in cytokine receptor signaling (19), we used quantitative mass spectrometry with tandem mass tags and antibody-based enrichment of tyrosine phosphorylated peptides. Global tyrosine phosphorylation in 32D-*Cbl*^KO^ cells expressing CBL WT or CBL C384Y, cultured in limiting IL3 (0.1ng/mL), was characterized to highlight the constitutively activated pathways in CBL C384Y cells. The global proteome as well as phosphoserine (pS) and phosphothreonine (pT) peptides were quantitated in parallel in the same samples (Supplementary Table 1-4). Among proteins detected in both global and phosphoproteomic datasets, we found no significant differences in the global proteome or proteome-normalized pS and pT peptides between cells expressing CBL WT and CBL C384Y. In contrast, we found that proteome-normalized tyrosine phosphorylation of CBL, LYN, and multiple proteins in the PI3K signaling pathway (e.g., PIK3R1/2, PIK3CB/D and PIK3AP1) were significantly increased in cells expressing CBL C384Y compared to CBL WT (Figure 1J). Phosphorylation of LYN, a SRC family kinase, on tyrosine 397 (pY397), was among the most significantly increased phosphotyrosine sites in CBL C384Y cells, and has been shown to stimulate LYN kinase activity (30). Changes in tyrosine phosphorylation that would indicate activation of the MAPK or JAK-STAT pathways were not observed in cells expressing CBL C384Y. Notably, MAPK1 pY185 and MAPK3 pY205, which indicate MAPK pathway activation, and STAT5A pY694 and STAT5B pY699, which indicates JAK activity, were significantly reduced in cells expressing mutant C384Y (Figure 1J). Thus, our mass spectrometry-based analysis of tyrosine phosphorylation implicated a CBL-LYN-PI3K axis in the enhanced cell proliferation associated with expression of mutant *CBL* in 32D cells.

To determine if LYN kinase activation was a common feature of *CBL* mutant expression, we measured LYN pY397 by western blot in cells expressing a series of mutant *CBL* alleles. In line with our phosphoproteomic results, LYN pY397 levels were increased in cells expressing *CBL* mutants compared to *CBL* WT (Figure 1K and Supplementary Figure 5). Total LYN protein was also increased in *CBL* mutant cell lines. Consistent with enhanced LYN activation, western blot analysis of 32D and TF1 cells expressing *CBL* mutants also revealed increased CBL phosphorylation at known LYN target sites Y700 and Y774 (25). Our global proteomic analysis also demonstrated a significant increase in tyrosine phosphorylation of PI3K-associated proteins, suggestive of increased signaling in the PI3K pathway, though the biological roles of the specific phosphotyrosine sites that were detected have not been reported. We therefore used western blot to measure AKT pS473 and ribosomal S6 pS235/236, which are directly indicative of signaling in the PI3K pathway. Indeed, phosphorylation of AKT and ribosomal S6 were increased in *CBL* mutant cells compared to *CBL* WT cells (Figure 1K and Supplementary Figure 5). AKT and S6 phosphorylation were unchanged in 32D-*Cbl*^KO^ cells even though LYN protein and phosphorylation were increased compared to 32D-*Cbl*^WT^ cells, highlighting that expression of CBL mutant protein was required for increased PI3K/AKT signaling. Altogether, these data indicate that gain-of-function *CBL* mutants are associated with increased LYN kinase activity, CBL phosphorylation, and activation of the PI3K/AKT signaling pathway.

### CBL mutants enhance cytokine signaling and cell proliferation via increased interaction with LYN

*CBL* mutations in myeloid malignancies disrupt the RING domain, and therefore E3 ubiquitin ligase function, while the adaptor domains, including the TKB domain, PRR and C-terminal tyrosine phosphorylation sites, remain intact (21). Thus, we hypothesized that the functions of CBL adaptor domains play a functional role in the increased cytokine signaling and cell proliferation in *CBL* mutant-expressing cells. To explore this possibility, we generated GFP-labeled double mutants comprising the C384Y RING domain mutation with secondary mutations in the TKB domain (G306E), PRR (Δ477-688), or C-terminal phosphotyrosine sites (Y700/731/774F) in 32D-*Cbl*^KO^ cells (Figure 2A). Single and double-mutant cells were mixed with dTomato-labeled *CBL* WT cells at a ratio of 1:10 and the GFP:dTomato ratio was measured by flow cytometry to assess for the relative rates of cell proliferation. The competitive advantage of cells expressing CBL C384Y was significantly reduced with mutation of the TKB domain, PRR, or phosphotyrosine residues, indicating that CBL’s adaptor domains are critical for the proliferative advantage of cells expressing mutant *CBL* (Figure 2B).

**Figure 2.**
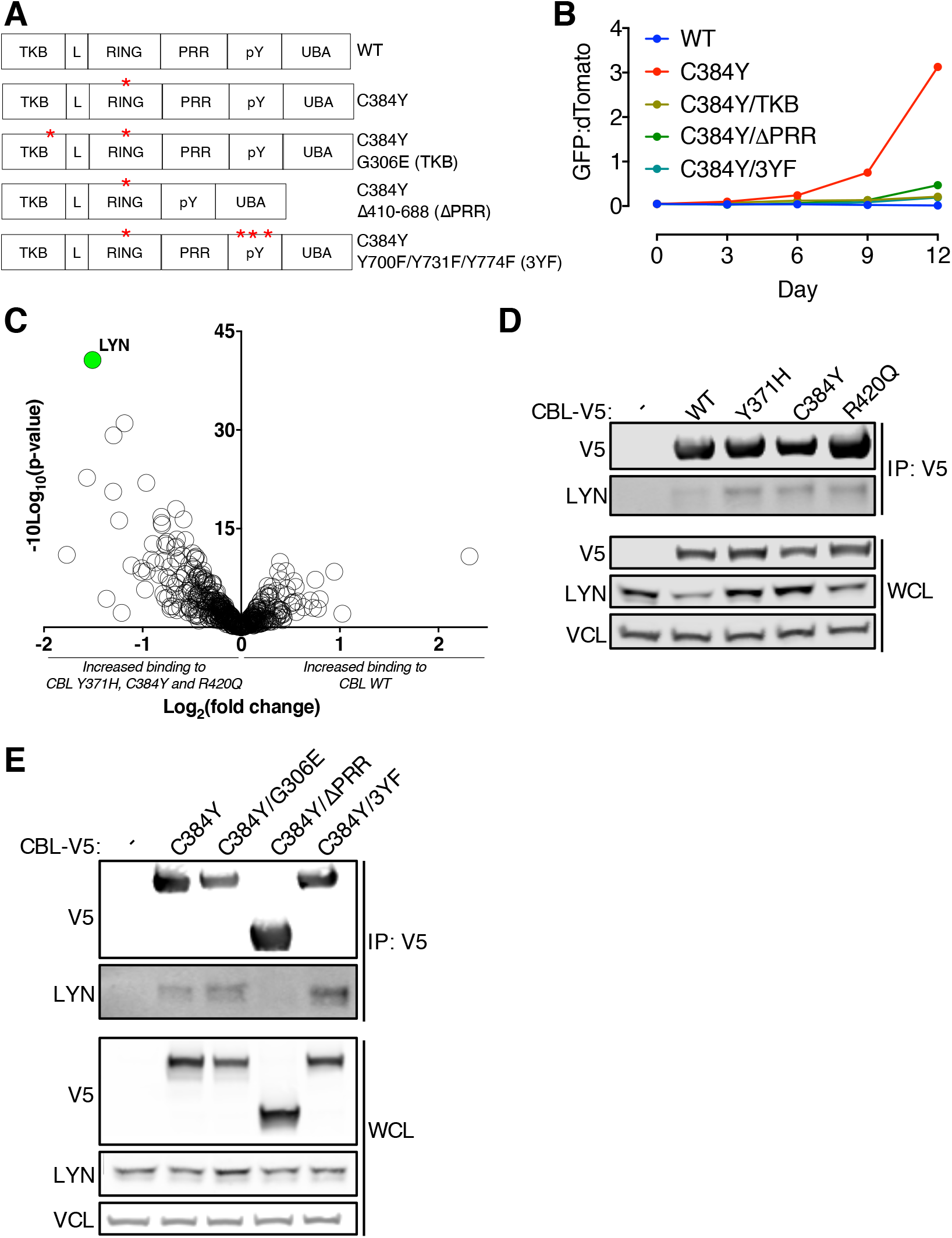
LYN binding to the proline-rich region of CBL is enhanced in cells expressing CBL mutants. (A) Schematic of CBL variants with mutation of the RING domain (C384Y) in combination with the tyrosine kinase binding (TKB) domain (G306E), proline-rich region (PRR) (**Δ**PRR), or C-terminal phosphotyrosine sites (Y700F/Y731F/Y774F). (B) Competition between CBL WT (dTomato) and CBL double mutants (GFP) expressed in 32D-*Cbl*^KO^ cells. (C) Mass spectrometry analysis of V5 immunoprecipitation (IP) samples from 32D-*Cbl*^KO^ cells expressing V5-tagged CBL WT, Y371H, C384Y or R420Q. Volcano plot showing log_10_ p-values and log_2_ fold change in protein abundance between CBL mutants (pooled data from Y371H, C384Y and R420Q) and CBL WT samples. (D) Western blot for V5, LYN and vinculin (VCL) in anti-V5 IP samples and whole-cell lysates (WCL) from 32D-*Cbl*^KO^ cells expressing V5-tagged CBL WT, Y371H, C384Y and R420Q. (E) Western blot for V5, LYN and vinculin (VCL) in anti-V5 IP samples and whole-cell lysates (WCL) from 32D-*Cbl*^KO^ cells expressing V5-tagged CBL WT, C384Y, C384Y/G306E, C384Y/**Δ**PRR, or C384Y/3YF.

Based on the key role of CBL adaptor domains in the proliferative advantage of cells expressing *CBL* mutants, we hypothesized that CBL adaptor domains promote increased cytokine sensitivity via specific protein-protein interactions with these domains. To address this possibility, we employed immunoprecipitation followed by mass spectrometry (IP-MS) to characterize the global CBL interactome in 32D-*Cbl*^KO^ cells expressing V5-tagged CBL WT or mutants Y371H, C384Y or R420Q (Supplementary Tables 5 and 6). Comparison of the WT and mutant CBL interactomes revealed significantly increased binding of LYN to CBL mutants compared to CBL WT (Figure 2C), and we confirmed this finding by IP followed by western blot (IP-WB) in both 32D and TF1 cell lines (Figures 2D and Supplementary Figure 6). To define the CBL adaptor domain(s) required for the interaction between mutant CBL and LYN, we performed additional IP-WB experiments using 32D-*Cbl*^KO^ cells expressing the CBL C384Y RING domain mutant with secondary mutations in the TKB domain, PRR or C-terminal phosphotyrosine sites. Secondary mutation of only the PRR abolished LYN binding to CBL C384Y, whereas mutations in the TKB domain or C-terminal phosphotyrosine sites had no effect (Figure 2E). Similar results were obtained in TF1 cells (Supplementary Figure 6). Overall, our global proteomic analyses demonstrated that enhanced proliferation of cells expressing *CBL* mutants was associated with increased LYN kinase activation, LYN protein levels, and interaction of LYN with mutant CBL.

### LYN interaction with mutant CBL increases PIK3R1 recruitment and downstream PI3K/AKT signaling

To explore the role of increased CBL-LYN interaction in *CBL* mutant cell lines, we sought to identify proteins with differential interaction with CBL in the context of LYN binding. In a nearest neighbors analysis of our CBL IP-MS data to identify proteins with a pattern of binding most similar to LYN, the PI3K signaling adaptor protein PIK3R1 was among the top hits. Moreover, tyrosine phosphorylation of PIK3R1 was markedly increased in CBL C384Y-expressing cells compared to CBL WT-expressing cells. Immunoprecipitation of V5-tagged CBL followed by western blot for PIK3R1 confirmed that CBL mutants Y371H, C384Y and R420Q bound significantly more PIK3R1 compared to CBL WT (Figure 3B). Consistent with previous studies (31, 32), CBL C384Y with a secondary Y731F mutation was unable to bind PIK3R1 in both 32D and TF1 cells, indicating that tyrosine phosphorylation at position 731 of mutant CBL was essential for the interaction with PIK3R1 (Figures 3C and D). Deletion of CBL’s proline-rich region, which was required for the CBL-LYN interaction, also led to a significant decrease in the CBL-PIK3R1 interaction (Figure 3C), suggesting a potential connection between the CBL-LYN interaction and increased PIK3R1 binding. Indeed, the interaction between PIK3R1 and V5-tagged CBL C384Y was also reduced in *Lyn* knockout 32D cells engineered by CRISPR-Cas9-mediated gene editing (Figure 3E). Thus, increased CBL-LYN interaction enhances the binding of the PI3K signaling adaptor PI3KR1 to CBL phosphotyrosine Y731.

**Figure 3.**
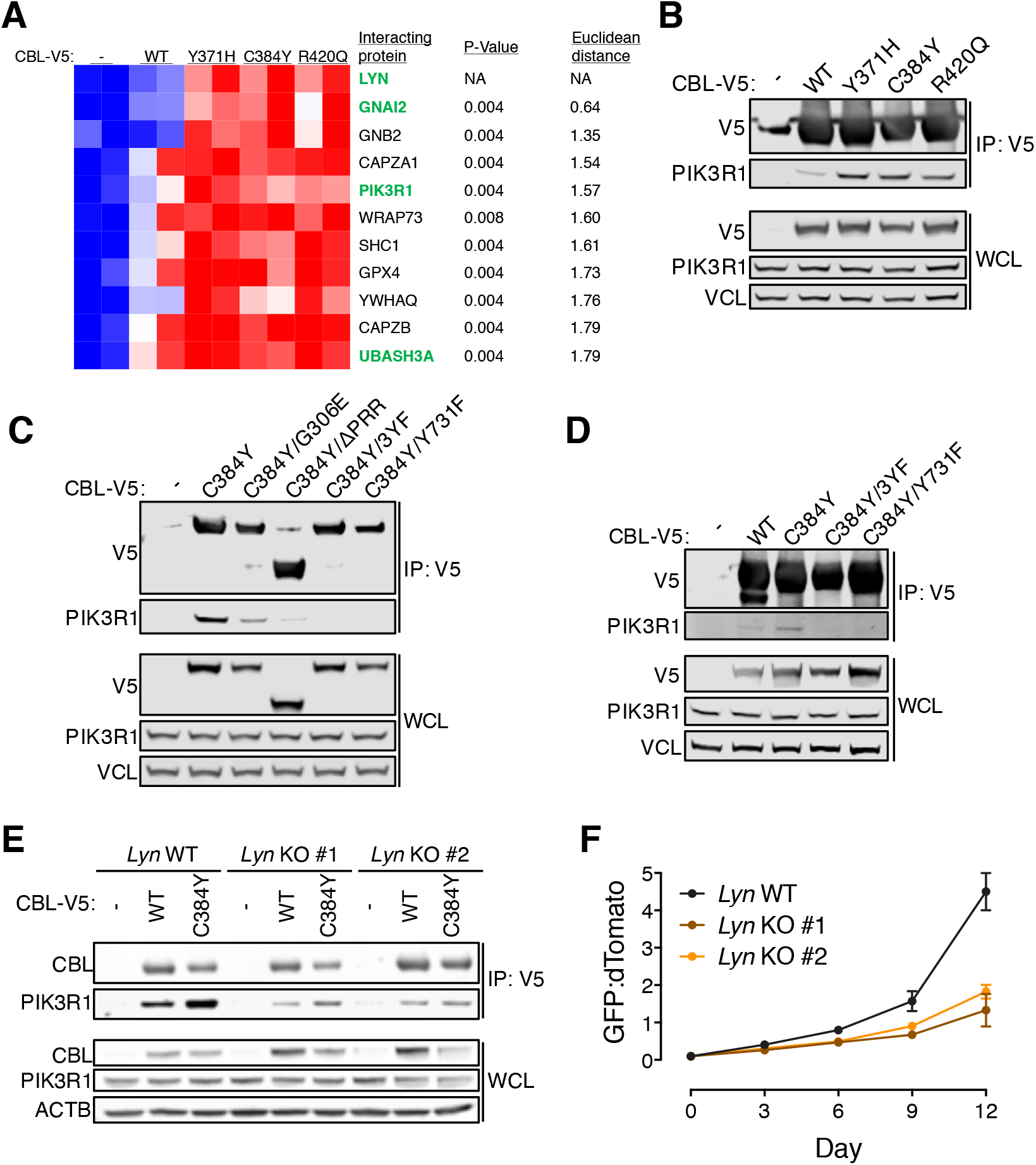
LYN facilitates CBL-PIK3R1 interaction and increased proliferation of CBL mutant cells. (A) Heat map depicting relative protein abundance in V5 immunoprecipitation (IP) samples analyzed by mass spectrometry. Top 10 proteins correlated with LYN by nearest neighbors analysis are shown. Proteins highlighted in green also showed significantly increased tyrosine phosphorylation in 32D cells expressing CBL C384Y compared to CBL WT. (B and C) Western blot for V5, PIK3R1 and vinculin (VCL) in anti-V5 IP samples and whole-cell lysates (WCL) from 32D-*Cbl*^KO^ cells expressing V5-tagged (B) CBL WT, Y371H, C384Y and R420Q or (C) CBL WT, C384Y, C384Y/G306E, C384Y/**Δ**PRR, C384Y/3YF or C384Y/Y731F. (D) Western blot for V5, PIK3R1 and VCL in anti-V5 IP samples and WCL from TF1-*CBL*^KO^ cells expressing V5-tagged CBL WT, C384Y, C384Y/3YF or C384Y/Y731F. (E) Western blot for V5, PIK3R1 and **β**actin (ACTB) in anti-V5 IP samples and WCL from 32D-*Lyn*^KO^ cells expressing V5-tagged CBL WT or C384Y. (F) Competition between CBL WT (dTomato) and C384Y (GFP) expressing 32D cells on *Lyn*^WT^ and *Lyn*^KO^ genetic backgrounds.

To test whether LYN was required for increased cell proliferation driven by CBL mutants, we compared the competitive advantage of CBL C384Y-expressing cells in *Lyn* WT and knockout 32D cells. In competition against CBL WT-expressing cells, the proliferative advantage of CBL C384Y-expressing cells was reduced by *Lyn* knockout (Figure 3F). In line with this result, *Lyn* knockout cells expressing CBL C384Y were at a competitive disadvantage when co-cultured with *Lyn* WT cells expressing CBL C384Y (Supplementary Figure 8). Moreover, *Lyn* knockout was associated with reduced phosphorylation of CBL and AKT in CBL C384Y-expressing cells (Supplementary Figure 9). Altogether, these experiments suggest that LYN drives proliferation of cells expressing mutant CBL via increased CBL phosphorylation, PIK3R1 recruitment and downstream PI3K/AKT signaling.

### Dasatinib inhibits LYN activation, PI3K signaling and proliferation in cells expressing *CBL* mutants

Given the central role of LYN in cytokine signaling downstream of CBL mutants, we hypothesized that cells expressing *CBL* mutants would be susceptible to pharmacologic inhibition of LYN. Dasatinib is a drug that inhibits ABL- and SRC-family kinases, including LYN (33). In proliferation assays, we found that the IC_50_ of dasatinib was lower in 32D and TF1 cells expressing CBL mutants compared to CBL WT (Figure 4A and Supplementary Figure 9A). Dasatinib also blocked the proliferative advantage of 32D cells expressing CBL Y371H, C384Y and R420Q in competition assays (Figures 4B-D). Similar results were observed in TF1 cells and 32D cells with CRISPR-Cas9-mediated editing of *Cbl* exon 8 splice sites (Supplementary Figures 9B-F). The inhibitory effect of dasatinib on proliferation of *CBL* mutant cell lines correlated directly with decreased phosphorylation of LYN, CBL, AKT and S6, as measured by western blot (Figure 4E and Supplementary Figure 10). Moreover, dasatinib treatment abrogated the interaction between mutant CBL and PIK3R1 (Figure 4F). Overall, these results demonstrated that dasatinib inhibited the effects of *CBL* mutants on cytokine signaling and cell proliferation.

**Figure 4.**
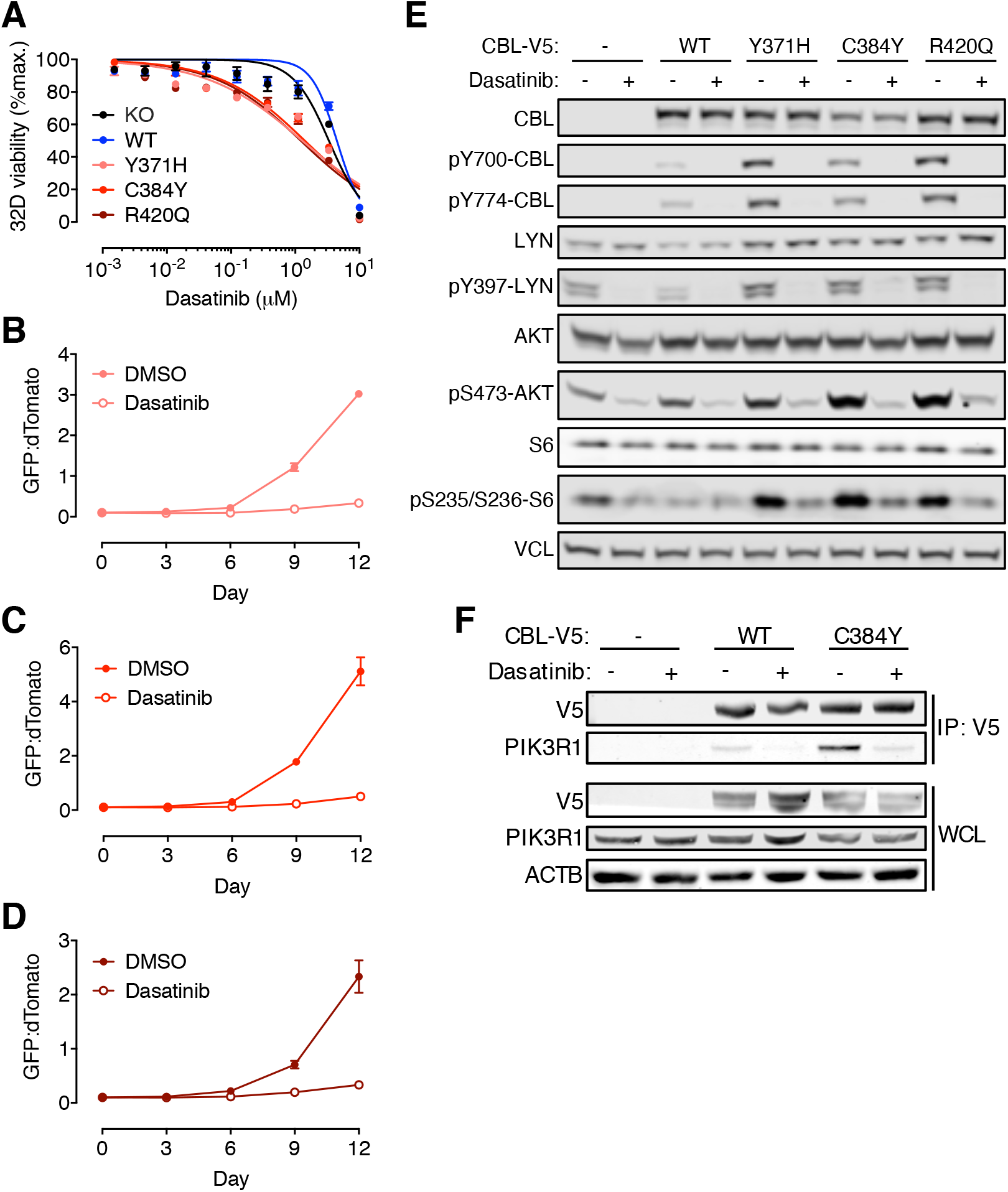
Dasatinib inhibits proliferation, CBL phosphorylation, PI3K/AKT signaling, and CBL-PIK3R1 interaction in *CBL* mutant cell lines. (A) Proliferation of 32D-*Cbl*^KO^ cells expressing luciferase (KO) or V5-tagged CBL WT, Y371H, C384Y, or R420Q in the presence of DMSO or a range of dasatinib concentrations over 3 days. (B-D) Competition between dTomato-labeled 32D-*Cbl*^KO^ cells expressing CBL WT and GFP-labeled 32D-*Cbl*^KO^ cells expressing (B) CBL Y371H, (C) CBL C384Y, or (D) CBL R420Q in the presence of DMSO (closed symbols) or 1**μ**M dasatinib (open symbols). (E) Western blot for total and phosphorylated CBL, LYN, AKT and S6 proteins in 32D-*Cbl*^KO^ cells expressing luciferase (-) or V5-tagged CBL WT, Y371H, C384Y, or R420Q treated with DMSO or 1**μ**M dasatinib for 2 hours. Western blot for vinculin (VCL) was used as a loading control. (F) Western blot for V5, PIK3R1 and **β**actin (ACTB) in anti-V5 IP samples and WCL from DMSO- or dasatinib-treated (1μM for 2 hours) 32D-*Cbl*^KO^ cells expressing luciferase (-) or V5-tagged CBL WT or C384Y.

The competitive advantage of *CBL* mutant cells was impaired but still detectable on a *Lyn* knockout genetic background (Figure 3F), indicating the presence of other contributors to the phenotype conferred by *CBL* gain-of-function mutations. To test whether these pathways involved another SRC-family kinase member, we assessed the effect of dasatinib on the proliferative advantage of 32D *Lyn* knockout cells expressing CBL C384Y. Remarkably, *Lyn* knockout rendered CBL C384Y-expressing cells completely insensitive to dasatinib treatment, implying that the reduction in proliferation of *CBL* mutant 32D cells treated with dasatinib was due to LYN inhibition (Supplementary Figure 11).

### Dasatinib inhibits clonogenicity and engraftment of *CBL*-mutated CMML patient samples *in vitro* and *in vivo*

We next sought to test the efficacy of dasatinib on *CBL*-mutant leukemic cells from patients with CMML. We assessed 4 *CBL* mutated patient samples in *in vitro* colony forming assays and *in vivo* xenograft studies (Figure 5A) (34). Up to 9 xenograft models were generated per patient and randomized to receive 14 days of 50mg/kg dasatinib or vehicle control 2-4 weeks after transplantation. Samples harbored *CBL* mutations in combination with a variety of other mutations frequently seen in CMML (Figure 5B and Supplementary Table 7). All samples were from patients with myeloproliferative features and splenomegaly consistent with the clinical presentation of *CBL* mutated CMML (Supplementary Table 8). Dasatinib reduced colony formation by approximately 50% compared to DMSO treatment in samples from all 4 patients (Figures 5C and D). For CMML samples #1, 2 and 4, dasatinib treatment of xenografted NSG-S mice was associated with decreased splenic disease burden as measured by the percentage of human CD45^+^ cells detected by flow cytometry and immunohistochemistry (Figures 5E and F). In mice xenografted with CMML sample #3, dasatinib was associated with a significant decrease in spleen weight and a trend towards decrease in the percentage of human CD45^+^ cells in the spleen and bone marrow, consistent with a reduction in absolute disease burden (Supplementary Figures 12A-D). Altogether, these experiments indicated that dasatinib effectively reduced the growth of primary, *CBL*- mutated CMML samples *in vitro* and *in vivo*.

**Figure 5.**
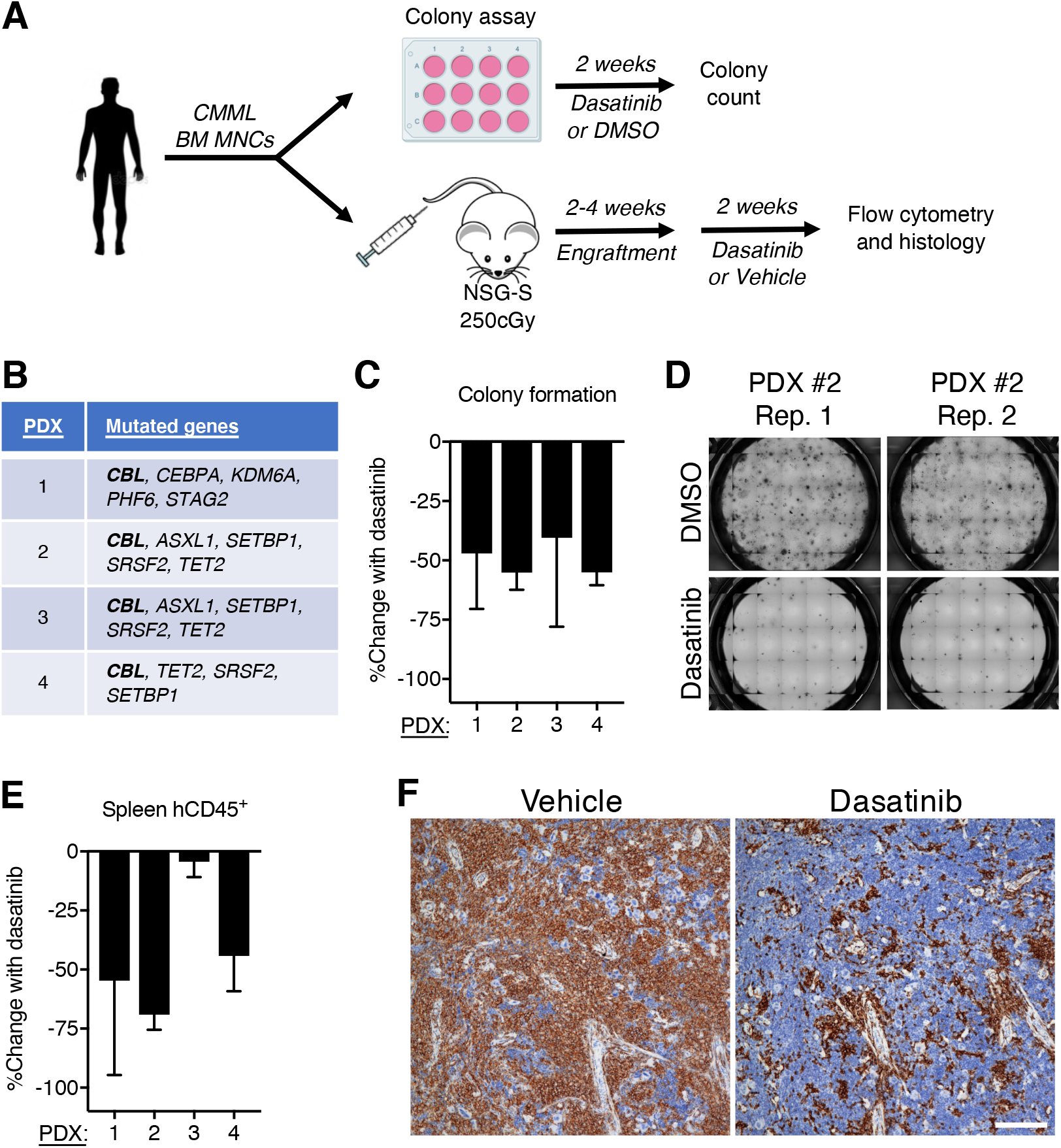
Dasatinib inhibits proliferation of *CBL*-mutated CMML patient samples *in vitro* and *in vivo*. (A) Schematic of the experimental workflow of CMML xenotransplantation model system. (B) Genes mutated in 4 CMML patient samples used for *in vitro* and *in vivo* experiments. (C) Percent change in colony formation with 250nM dasatinib compared to DMSO after 2 weeks of treatment (mean and S.D. of 3-4 replicate wells are depicted). (D) Representative images of colony formation by CMML patient sample #2 after 2 weeks in the presence of DMSO or 250nM dasatinib. (E) Percent change in human CD45^+^ cells in the spleens of NSG-S mice engrafted with CMML patient samples and treated with vehicle or 50mg/kg dasatinib via oral gavage over 2 weeks (mean and S.D. of 3-5 mice per group are depicted). (F) Anti-hCD45 immunohistochemisty in spleen tissue sections from mice engrafted with CMML patient sample #2 and treated with vehicle or dasatinib for 14 days.

## Discussion

We used phosphoproteomics, IP-MS, and functional analyses to identify signaling pathways that drive proliferation in a panel of cell lines expressing an allelic series of *CBL* mutations. This unbiased approach highlighted a mechanism whereby increased LYN activity and binding to mutant CBL enhances CBL tyrosine phosphorylation, PIK3R1 recruitment and downstream PI3K/AKT activation. In addition to defining this molecular pathway, our findings point to the therapeutic efficacy of LYN inhibition by dasatinib, which we validated in both cell lines and primary leukemia samples *in vitro* and *in vivo*.

*CBL* mutations in human myeloid malignancies selectively affect the protein’s E3 ubiquitin ligase function, suggesting that CBL’s role as a signaling adaptor is critical to the gain-of-function conferred by the mutant protein (21). Indeed, a prior study reported that the adaptor domains of mutant CBL were essential for increased signaling and cell proliferation (35), though the molecular basis for this finding was not fully addressed. Using IP-MS as an unbiased discovery tool, we identified and characterized the interaction between CBL and LYN (25–27) as an important component of oncogenic signaling driven by the proline-rich region of mutant CBL. We found an increase in total and activated LYN in both *CBL* knockout and gain-of-function mutant cell lines, suggesting that CBL’s E3 ubiquitin ligase function plays a role in regulation of LYN protein levels. We provide several lines of evidence that gain-of-function *CBL* mutations, unlike *CBL* knockout, specifically exploit LYN kinase activity to enhance downstream PI3K/AKT signaling.

Unlike other studies that observed hyperactivation of MAPK, JAK-STAT and PI3K/AKT pathways (2,6,15,23,24,28,35–41), our global characterization of tyrosine phosphorylation in cells expressing mutant *CBL* revealed selective activation of LYN and the PI3K/AKT pathway; in fact, tyrosine phosphorylation sites that directly indicate activation of the MAPK or STAT5 pathways were lower in our *CBL* mutant cells by MS analysis. There are several potential explanations for this discrepancy. First, most previous studies measured phospho-signaling using starve-stimulation assays, which provide an estimate of phosphorylation kinetics upon acute cytokine exposure (2,6,15,23,28,35–41). Here we performed an unbiased assessment of tyrosine phosphorylation during chronic cytokine exposure in order to identify signaling pathways that were activated and targetable in the steady state. Second, the effects of *CBL* mutations may depend on the cellular context. Along these lines, hematopoietic stem and progenitor cells from *Cbl* mutant mice showed increased PI3K/AKT, MAP kinase and STAT5 signaling, though there were differences in the signaling effects depending on the subpopulation analyzed (24). Third, two studies evaluated the effects of CBL mutants on a *Cbl^-/-^*/*Cblb^-/-^* genetic background (35, 41), which has the potential to augment or alter the signaling phenotype conferred by CBL mutant protein due to functional redundancy between *Cbl* and *Cblb* in mice (42–44). In patients with myeloid malignancies, *CBLB* mutations are uncommon and co-mutation of *CBL* and *CBLB* has not been described, suggesting that recurrent mutations in *CBLB* do not contribute significantly to oncogenesis. Though it is possible that *CBLB* modulates cytokine signaling in our cell lines models, we performed our analyses of *CBL* mutations in *CBLB* WT cells to accurately recapitulate human disease genetics. Fourth, the diversity of signaling adaptors and kinases associated with different cytokine receptors may determine which signaling pathways are affected by *CBL* mutations. Variability in the magnitude of the effect and pathway specificity could reflect differences in the role of mutant CBL downstream of FLT3, KIT, MPL, and receptors utilizing the common beta chain CD131 (i.e., receptors for IL3, IL5, and GM-CSF) (45, 46).

The SRC-family kinase inhibitor dasatinib effectively reduced proliferation and phospho-signaling in our *CBL* mutant cell lines, consistent with previous observations (28,39,40,47); and we provide the first evidence that dasatinib inhibits the proliferation of primary, human CMML samples *in vitro* and *in vivo*. The anti-proliferative effects of *in vivo* dasatinib treatment were relatively consistent across 4 different *CBL*-mutated CMML samples that were derived from patients with myeloproliferative disease with splenomegaly. Each patient sample had a unique combination of genes that were co-mutated with *CBL*, including 2 samples with *ASXL1* mutations, which have been associated with reduced overall survival in multiple studies (16,48–50). In our *in vivo* studies, mice engrafted with patient-derived CMML cells were only treated with dasatinib once daily for 10 days prior to analysis. Increasing the treatment period and frequency as well as the dose of dasatinib may lead to improved responses.

The effects of *CBL* mutant expression were significantly reduced but not abolished in *Lyn* knockout cells, indicating the presence of additional protein-protein interactions and signaling pathways involved in the gain-of-function conferred by *CBL* mutations; this result also highlights the likelihood that effective treatment will require combination therapy targeting multiple pathways (24,51,52). Molecular characterization of LYN-independent signaling pathways in our model systems will reveal additional therapeutic strategies for patients with *CBL*-mutated myeloid disease.

Somatic mutations in *CBL* are typically detected in subclones that arise during disease progression (53); *CBL* mutations can also function as disease-initiating mutations, consistent with their presence in a subset of healthy individuals with clonal hematopoiesis (54–56). We observed that dasatinib inhibited the proliferation of human CMML samples with either subclonal or clonal *CBL* mutations. This may indicate that *CBL* mutant cells contribute disproportionately to colony formation *in vitro* or leukemia expansion in NSG-S mice, which ultimately selects for a dasatinib-sensitive cell population. Alternatively, dasatinib may have broadly inhibitory effects on human CMML independently of the presence of *CBL* mutations. Ongoing studies aimed at determining whether the effects of dasatinib are correlated with particular CMML mutational profiles may guide the translational application of our findings.

## Materials and Methods

### Analysis of *CBL* mutations in patient samples

Patients undergoing evaluation for various hematologic disorders (2015–2019) were consented for targeted gene sequencing using a panel developed at the Brigham and Women’s Hospital and Dana-Farber Cancer Institute (29). The frequency and type of *CBL* mutations detected in 191 de-identified patients were catalogued.

### Cell culture

32D cells (ATCC CRL-11346) and BaF3 cells (DSMZ ACC 300) were maintained in RPMI 1640 (Corning, 10-040-CV) with 10% fetal bovine serum (FBS, Omega Scientific, 91356), 1x penicillin-streptomycin-glutamine supplement (ThermoFisher Scientific, 10378016), and 5ng/ml recombinant murine IL-3 (PeproTech, 213-13). TF1 cells (ATCC CRL-2003) were maintained in RPMI 1640 with 10% fetal bovine serum, 1x penicillin-streptomycin-glutamine supplement, and 5ng/ml recombinant human GM-CSF (Miltenyi Biotec, 130-093-866). HEK293T cells (ATCC CRL-3216) were maintained in DMEM (Corning, 10-013-CV) with 10% fetal bovine serum. All cells were culture at 37°C/5%CO_2_.

### Cloning and lentiviral packaging

For sgRNA constructs, annealed oligonucleotides were phosphorylated with T4 polynucleotide kinase (NEB, M0201S) and cloned into BsmBI-linearized, gel purified LentiCRISPRv2 (Cas9 + sgRNA for *Cbl* and *Lyn* knockouts), sgRNA.SFFV.TagBFP (sgRNA only for non-targeting or *Cbl* exon 1 targeting), or sgRNA.SFFV.RFP657 (sgRNA only for *Cbl* intron 7/8 targeting) using T4 DNA ligase (NEB, M0202S) (57). Donor constructs for *CBL* mutants were generated via site-directed mutagenesis (NEB Q5 Site-Directed Mutagenesis Kit, E0554S) of pDONR221 plasmid containing human *CBL* cDNA (NM_005188.4) with a C-terminal V5 tag. Expression constructs for *CBL* mutants were generated via Gateway LR Clonase (ThermoFisher Scientific, 11791100) reaction between pDONR221 plasmid (ThermoFisher Scientific, 12536017) containing V5-tagged wild-type or mutant *CBL* cDNA and lentiviral destination plasmid pRRL-SFFV-IRES-GFP or -dTomato (kindly provided by Christopher Baum and Axel Schambach, Hannover Medical School, Hannover, Germany). All constructs were confirmed by Sanger sequencing (Eton Biosciences, Boston, MA) and alignment using Benchling Biology Software (2018, https://benchling.com). Oligonucleotides (Eton Bioscience, Boston, MA) used for generating sgRNA constructs and *CBL* cDNA mutagenesis are listed below:

#### Oligonucleotides for sgRNAs

*Cbl* exon 1 targeting (*Cbl* KO):

FWD: CAC CGT CTT GTC CAC CGT GCA GGG A

REV: AAA CTC CCT GCA CGG TGG ACA AGA C

*Cbl* intron 7-1 targeting (*Cbl* exon 8 mis-splicing):

FWD: CAC CGT ATA ATT CAT ATT GTT CCT T

REV: AAA CAA GGA ACA ATA TGA ATT ATA C

*Cbl* intron 7-2 targeting (*Cbl* exon 8 mis-splicing):

FWD: CAC CGT CTC ATT TTT CTT TCA CCA A

REV: AAA CTT GGT GAA AGA AAA ATG AGA C

*Cbl* intron 8-1 targeting (*Cbl* exon 8 mis-splicing):

FWD: CAC CGG CCT CAC GTC GTG GCA GGT A

REV: AAA CTA CCT GCC ACG ACG TGA GGC C

*CBL* exon 1 targeting (*CBL* KO):

FWD: CAC CGC AAC GTG AAG AAG AGC TCT G

REV: AAA CCA GAG CTC TTC TTC ACG TTG C

*Lyn* exon 4 targeting (*Lyn* KO #1):

FWD: CAC CGA GAG GAA CAA GGT GAC ATT G

REV: AAA CCA ATG TCA CCT TGT TCC TCT C

*Lyn* exon 7 targeting (*Lyn* KO #2):

FWD: CAC CGC CAA CTT AAT GGA CTC CCG G

REV: AAA CCC GGG AGT CCA TTA AGT TGG C

#### Oligonucleotides for CBL cDNA site-directed mutagenesis

*CBL* c.1111T>C (Y371H):

FWD: ATA TGA ATT ACA CTG TGA GAT GGG

REV: TGT TCC TGG GTC ACT TTG

*CBL* c.1151G>A (C384Y):

FWD: TGT AAA ATA TAT GCT GAA AAT GAT AAG G

REV: TAG TTG GAA TGT GGA GCC

*CBL* c.1259G>A (R420Q):

FWD: CCT TTC TGC CAA TGT GAA ATT AAA GG

REV: ACA GCC CTG ACC TTC TGA

*CBL* c.917G>A (G306E, TKB domain mutation):

FWD: TGG GCT ATT GAG TAT GTT ACT GC

REV: CTG ACC CAG ACG AGT ACA G

*CBL* c.1228-2064del (Δ477-688, PRR deletion):

FWD: GGG GAG CAA TGT GAG GGT

REV: GGC ACC AGC CAA TTC CTT C

*CBL* c.2099A>T (Y700F, phosphotyrosine mutation):

FWD: GAC ACA GAG TTC ATG ACT CCC

REV: CTC TTC ACC CTC ACA TTG

*CBL* c.2192A>T (Y731F, phosphotyrosine mutation):

FWD: AGC TGT ACG TTT GAA GCA ATG TAT AAT ATT C

REV: ATC AAT CTG CTG GTC GCA ATC

*CBL* c.2321A>T (Y774F, phosphotyrosine mutation):

FWD: GAT GAT GGG TTT GAT GTC CCA AAG

REV: CTC ATT TTC TGA CTC CTC G

Lentivirus was produced in a 6cm^2^ dish by transient transfection of HEK293T cells using 18μL TransIT-LT1 (Mirus, MIR2304) in 150-170μL Opti-MEM reduced serum medium (ThermoFisher Scientific, 31985088) containing 3μg lentiviral expression plasmid, 4μg psPAX2 packaging plasmid (Addgene, 12260), and 1.5μg pCMV-VSV-G envelope plasmid (Addgene, 8454). Lentiviral supernatant was harvested at 48 hours post-transfection and passed through a 0.45μm syringe filter (Pall, 4614) prior to use for cell transduction.

### Cell line generation

For lentiviral transduction, cells were plated in 48-well plates (2×10^5^ cells/well in 600μL) and mixed with polybrene (Santa Cruz Biotechnology, 134220) to a final concentration of 4μg/mL and 100-200μL undiluted lentiviral supernatant. Plates were centrifuged at 1050 x g at 37°C in an Eppendorf 5910R centrifuge (Hauppauge, NY) for 1 hour and subsequently cultured overnight at 37°C/5%CO_2_. Cells were harvested, washed twice in phosphate buffer saline (PBS) (Corning, 21-040-CV), re-plated in 6-well plates with RPMI-based media containing the appropriate cytokine, and expanded at 37°C/5%CO_2_. For LentiCRISPRv2 and lentiCas9-Blast (Addgene, 52962) transductions, selection with puromycin (ThermoFisher Scientific, A1113802, 2μg/mL final concentration) and blasticidin (ThermoFisher Scientific, A1113902, 10μg/mL final concentration), respectively, was started on day 2 post-transduction. Cells transduced with lentiviral constructs containing fluorescent proteins were sorted using a Sony SH800S or MA900 cell sorter (Tokyo, Japan).

### PCR amplification and sequencing of *Cbl* exons 7-9

Total RNA was extracted from 32D-Cas9 cell lines expressing sgRNAs targeting *Cbl* introns 7 and 8 using an RNeasy Mini Kit (QIAGEN, 74106). cDNA was prepared using a QuantiTect Reverse Transcription Kit (QIAGEN, 205313). cDNA was subsequently used for PCR amplification of a 382bp fragment, including the splicing junctions between *Cbl* exons 7-9, using Q5 High-Fidelity Master Mix (NEB, M0492) and the following primers: (FWD) ATT TGT TTC CTG ATG GAC GAA ATC; (REV) GGC CTT TCC ACC TTG GCA. PCR cycles were performed using an Eppendorf Mastercycler (Hauppauge, NY) as follows: 30 seconds at 98°C; 25 x (10 seconds at 98°C, 10 seconds at 64°C, 10 seconds at 72°C); 2 minutes at 72°C; hold at 10°C. PCR amplicons were separated by agarose gel electrophoresis, individually isolated using a Gel Extraction Kit (QIAGEN, 28704), analyzed by Sanger sequencing (Eton Bioscience, Boston, MA), and aligned using Benchling Biology Software (2018, https://benchling.com).

### Cytokine dose response assays

32D, BaF3 or TF1 cells were harvested from culture and counted using a ViCELL-XR (Beckman Coulter, Indianapolis, IN). Cells were washed 3 times in PBS to remove remaining cytokine. 10^4^ cells were plated in a 96-well flat bottom plate in 200μL RPMI containing 10% FBS and 1x penicillin-streptomycin-glutamine supplement. A Tecan D300e Digital Dispenser (San Jose, CA) was used to dispense the appropriate amount of recombinant mouse IL-3 (for 32D and BaF3) or recombinant human GM-CSF (for TF1) in dH_2_O + 0.1% Triton X-100 as per the manufacturer’s instructions. Each cytokine concentration was tested in triplicate. CellTiter-Glo (Promega, G7570) was used to measure relative cell viability as per the manufacturer’s instructions at 72 hours. An EnVision 2105 plate reader (Perkin Elmer, Shelton, CT) was used to measure luminescence. Luminescence was normalized to the highest cytokine concentration for each cell line. Non-linear curves were fitted to the data and EC_50_ values were calculated using GraphPad Prism 7 (San Diego, CA).

### Cell competition assays

Cells expressing different fluorescent proteins were harvested from culture and counted using a ViCELL-XR (Beckman Coulter, Indianapolis, IN). Cells were washed 3 times in PBS to remove remaining cytokine. Cells were resuspended in RPMI containing 10% fetal bovine serum and penicillin-streptomycin-glutamine supplement with 0.1ng/mL recombinant mouse IL-3 (for 32D and BaF3) or 0.5ng/mL recombinant human GM-CSF (for TF1), mixed at a 10:1 ratio (final cell concentration 5×10^4^ cells/mL), and plated in triplicate in a 96-well flat bottom plate (200μL/well). Every 3-4 days, 20μL cell suspension was passed into a new 96-well plate with 180μL fresh media and the ratio of cells expressing different fluorescent proteins was measured by flow cytometry using a BD FACSCanto II (BD Biosciences, San Jose, CA). For competition assays with dasatinib (Selleck Chemicals, S7782), a Tecan D300e digital dispenser (San Jose, CA) was used to dispense the appropriate volume of dasatinib in DMSO as per the manufacturer’s instructions. Wells were DMSO-normalized to maintain the same concentration of DMSO across varying dasatinib concentrations.

### Quantitative Proteomics and Phosphoproteomics

#### Proteome and phosphoproteome analyses by liquid chromatography-tandem mass spectrometry (LC-MS/MS)

The experimental setup for quantitative proteomics and phosphoproteomics is presented in Supplementary Table 1. Cell pellets were lysed for 30 minutes with ice cold buffer containing 8M urea, 75mM NaCl, 50mM Tris-HCl (pH 8.0), 1mM EDTA, 2μg/mL aprotinin (Millipore Sigma, A1153), 10μg/mL leupeptin (Millipore Sigma, 11017101001), 1mM phenylmethylsulfonyl fluoride (Millipore Sigma, P7626), Phosphatase Inhibitor Cocktail 2 (1:100, vol/vol), Phosphatase Inhibitor Cocktail 3 (1:100, vol/vol), and 10mM NaF. Samples were spun at 20,000 x *g* for 10 minutes and a BCA assay (ThermoFisher Scientific, 23225) was used to determine the concentration of protein in each cleared lysate. Lysis buffer was used to equalize the protein concentration of each lysate to 8mg/mL. Protein disulfide bonds were reduced with 5mM dithiothreitol (ThermoFisher Scientific, R0861) at 25°C for 45 minutes. The proteins were alkylated in the dark using 10mM iodoacetamide at 25°C for 45 minutes. Lysates were then diluted 1:4 using 50mM Tris-HCl, pH 8.0, to lower the urea concentration to 2M. LysC (Wako, 125-05061) was added to each lysate at a 1:50 enzyme-to-substrate ratio and samples were digested at 25°C for 2 hours. Following LysC digestion, trypsin (Promega, V5280) was added at a 1:50 enzyme-to-substrate ratio and the samples were digested at 25°C overnight. The digestion was quenched with 100% formic acid (FA) to reach a volumetric concentration of 1% FA. Samples were spun at 5000 x *g* for 5 minutes to remove precipitated urea and then desalted using Sep-Pak C18 columns (Waters, 500mg WAT043395). Sep-Pak columns were conditioned with 1 × 5mL 100% acetonitrile (ACN), 1 × 5mL 50% ACN/0.1% formic acid (FA), and 4 × 5mL 0.1% trifluoroacetic acid (TFA). Each sample was loaded onto a column and washed with 3 × 5mL 0.1% TFA and 1 × 5mL 1% FA. Peptides were eluted off the column with 2 × 3mL 50% ACN/0.1% FA and dried down. Peptides were resuspended in 3% ACN, 0.1% FA and a BCA assay was used to determine the concentration of peptide in each sample.

Desalted peptides from each sample (500μg) were labeled with 10-plex Tandem Mass Tag (TMT) reagents (ThermoFisher Scientific, 90110) (58). A 5% aliquot of each sample was used for global proteomic analysis, dried in a Speed-Vac, and resuspended in 3% ACN/0.1% formic acid prior to LC-MS/MS analysis. The remaining sample was utilized for phosphopeptide enrichment. Briefly, for enrichment of phosphotyrosine (pY) peptides, the combined TMT10-labeled sample was resuspended in 2mL of IAP buffer (50mM MOPS, pH 7.2, 10mM sodium phosphate, and 50mM NaCl). The sample was mixed with 100ug of pY antibody (Cell Signaling Technology, 8954) and incubated at 4°C for 1 hour with end over end rotation. Prior to the incubation, pY antibody was washed 3x in IAP buffer. Following the incubation, enriched peptides were washed 4x in PBS, and eluted from the beads 2x with 0.15% TFA. Eluted peptides were desalted on the C18 StageTips. Briefly, StageTips were calibrated with 100% MeOH, 50% MeCN/0.1% FA, and with 0.1% FA. Peptides were eluted from the StageTips with 50% MeCN/ 0.1% FA, dried to completion, and resuspended in 3% ACN/0.1% formic acid prior to ESI-LC-MS/MS analysis. For enrichment of phosphoserine (pS) and phosphothreonine (pT) peptides, Ni-NTA agarose beads were used to prepare Fe^3+^- NTA agarose beads, and peptides were reconstituted in 80% ACN/0.1% TFA and incubated with 10μL of the Fe^3+^-IMAC beads for 30 minutes. Samples were then centrifuged, and the supernatant containing unbound peptides was removed. The beads were washed twice and then transferred onto equilibrated C18 StageTips with 80% ACN/0.1% TFA. StageTips were rinsed twice with 1% FA and eluted from the Fe^3+^- IMAC beads onto C18 StageTips with 70μL of 500mM dibasic potassium phosphate, pH 7.0, a total of three times. C18 StageTips were then washed twice with 1% FA, followed by elution of the phosphopeptides from the C18 StageTips with 50% ACN/0.1% FA twice. Samples were dried down and resuspended in 3% ACN/0.1% FA prior to LC-MS/MS analysis.

Global proteome and phosphoproteome samples were analyzed on an Orbitrap Fusion Lumos Mass Spectrometer (ThermoFisher Scientific). Peptides were separated on an Easy nLC 1200 UHPLC system (ThermoFisher Scientific) using an in-house packed 20cm x 75μm diameter C18 column (1.9μm Reprosil-Pur C18-AQ beads; Picofrit 10μm opening, Dr. Maisch GmbH (New Objective)). The column was heated to 50°C using a column heater (Phoenix-ST). The flow rate was 0.2μl/min with 0.1% FA and 2% ACN in water (A) and 0.1% FA, 90% ACN (B). The peptides were separated with a 6%–30% B gradient in 84 minutes. Parameters were as follows: MS1: resolution – 60,000, mass range – 350 to 1800m/z, AGC Target 4.0e^5^, Max IT – 50ms, charge state include - 2-6, dynamic exclusion – 5s, top 20 ions selected for MS2; MS2: resolution – 50,000, high-energy collision dissociation activation energy (HCD) – 38, isolation width (m/z) – 0.7, AGC Target – 1.0e^5^, Max IT – 250ms.

#### CBL Interaction Proteomics

The experimental setup for quantitative CBL interaction proteomics is presented in Supplementary Table 2. For immunoprecipitation samples, beads were washed once with cell lysis buffer, 3x with PBS, and subsequently resuspended in 90μL digestion buffer (2M Urea, 50 mM Tris-HCl) with 2μg trypsin. Samples were incubated at 25°C for 1 hour with shaking at 700rpm. The supernatant was removed and placed in a fresh tube. The beads were then washed twice with 50μL digestion buffer. The combined supernatants were reduced (2μL 500mM DTT for 30 minutes at 25°C), alkylated (4μL 500mM IAA for 45 minutes in the dark) and digested overnight with an additional 2μg trypsin. Samples were then quenched with 20μL 10% FA, desalted on 10mg Oasis cartridges, and labeled with 10-plex TMT reagents.

For immunoprecipitation samples, reconstituted peptides were separated using an online nanoflow EASY-nLC 1000 UHPLC system (ThermoFisher Scientific) and analyzed on a benchtop Orbitrap Q Exactive plus mass spectrometer (ThermoFisher Scientific). The peptide samples were injected onto a capillary column (Picofrit with 10μm tip opening / 75μm diameter, New Objective, PF360–75-10-N-5) packed in-house with 20cm C18 silica material (1.9 μm ReproSil-Pur C18-AQ medium, Dr. Maisch GmbH) and heated to 50°C in column heater sleeves (Phoenix-ST) to reduce backpressure during UHPLC separation. Injected peptides were separated at a flow rate of 200nL/min with a linear 230 minute gradient from 100% solvent A (3% ACN, 0.1% FA) to 30% solvent B (90% ACN, 0.1% FA), followed by a linear 9 minute gradient from 30% solvent B to 60% solvent B and a 1 minute ramp to 90% solvent B. Each sample was run for 260 minutes, including sample loading and column equilibration times. The Q Exactive instrument was operated in the data-dependent mode acquiring HCD MS/MS scans (R=17,500) after each MS1 scan (R=70,000) on the 12 top most abundant ions using an MS1 ion target of 3× 10^6^ ions and an MS2 target of 5×10^4^ ions. The maximum ion time utilized for the MS/MS scans was 120ms; the HCD-normalized collision energy was set to 27; the dynamic exclusion time was set to 20s, and the peptide match and isotope exclusion functions were enabled.

#### Data Analysis

All mass spectra were processed using the Spectrum Mill software package v6.0 pre-release (Agilent Technologies), which includes modules for TMT-based quantification. For peptide identification MS/MS spectra were searched against the mouse UniProt database to which a set of common laboratory contaminant proteins was appended. Search parameters included: ESI-Q Exactive-HCD scoring parameters, trypsin enzyme specificity with a maximum of two missed cleavages, 30% minimum matched peak intensity, +/−20ppm precursor mass tolerance, +/−20ppm product mass tolerance, and carbamidomethylation of cysteines and TMT labeling of lysines and peptide N-termini as fixed modifications. Allowed variable modifications for proteome data were oxidation of methionine, N-terminal acetylation, with a precursor MH+ shift range of 0 to 70 Da. Allowed variable modifications for pY data were oxidation of methionine, N-terminal acetylation, and pY, pS, or pT with a precursor MH+ shift range of −18 to 272 Da. For CBL interaction data, allowed variable modifications were oxidation of methionine, N-terminal acetylation, with a precursor MH+ shift range of 0 to 70 Da. Identities interpreted for individual spectra were automatically designated as valid by optimizing score and delta rank1-rank2 score thresholds separately for each precursor charge state in each LC-MS/MS while allowing a maximum target-decoy-based false-discovery rate (FDR) of 1.0% at the spectrum level.

### Western blot

Cells were lysed in buffer containing 150mM NaCl, 50mM Tris (pH 7.5), 1% NP40 (Millipore, 492018), and Halt protease/phosphatase inhibitor cocktail (ThermoFisher Scientific, 78446) for 15 minutes on ice. Protein lysates were pre-cleared by centrifuging at 21000 x g for 10 min at 4°C. Supernatants were harvested and the protein concentration was measured using the Pierce BCA protein assay kit (ThermoFisher Scientific, 23225). Samples were mixed with NuPAGE LDS sample buffer (ThermoFisher Scientific, NP0007) and reducing agent (ThermoFisher Scientific, NP0004) and boiled at 70°C for 10 minutes. Proteins were resolved by SDS-PAGE using NuPAGE 4-12% Bis-Tris protein gels (ThermoFisher Scientific, NP0336), and transferred to 0.45μm nitrocellulose membranes (Life Technologies, LC2001) by electrophoresis. Membranes were blocked in Odyssey blocking buffer (Li-Cor, 927-50000) for 1 hour at room temperature and subsequently incubated with primary antibodies in Odyssey blocking buffer overnight at 4°C. Membranes were washed 3 times in tris-buffered saline with 0.1% Tween 20 (TBS-T, Cell Signaling, 9997) at room temperature and subsequently incubated with secondary antibodies for 1 hour at room temperature in Odyssey blocking buffer. Membranes were washed 3 times in TBS-T at room temperature and visualized using a Li-Cor Odyssey CLx (Lincoln, NE). Immunoblot quantitation in Supplementary Figure 8 was performed using ImageJ software (59).

### Immunoprecipitation

Protein lysates were prepared as described above, an aliquot was saved for whole cell lysate analysis, and 1-2mg total protein lysate in 500μL was incubated with 50ul anti-V5 magnetic bead slurry (MBL International, M167-11) overnight with rotation at 4°C. Beads were separated by magnet and washed once with lysis buffer and twice with wash buffer (150mM NaCl, 50mM Tris pH 7.5, 1x Halt protease/phosphatase inhibitor cocktail). Samples were eluted from magnetic beads by boiling in LDS NuPAGE sample buffer with reducing agent at 70°C for 10 minutes. Proteins were resolved by SDS-PAGE and analyzed by immunoblot or mass spectrometry as described above.

### Antibodies

#### Primary antibodies

| **Antibody** | **Supplier** | **Catalog #** | **Dilution Used** |
| --- | --- | --- | --- |
| Anti-AKT (pan) | Cell Signaling Technologies | 4685 | 1:1000 |
| Anti-pS473-AKT | Cell Signaling Technologies | 4060 | 1:1000 |
| Anti-β-Actin | Cell Signaling Technologies | 3700 | 1:10000 |
| Anti-c-CBL | Cell Signaling Technologies | 2747 | 1:1000 |
| Anti-pY700-CBL | Abcam | ab76002 | 1:1000 |
| Anti-pY731-CBL | Abcam | ab52855 | 1:1000 |
| Anti-pY774-CBL | Abcam | ab32446 | 1:1000 |
| Anti-LYN | Cell Signaling Technologies | 2796 | 1:1000 |
| Anti-p85 | Cell Signaling Technologies | 4257 | 1:1000 |
| Anti-SRC family (phospho-Y418) | Abcam | ab40660 | 1:1000 |
| Anti-S6 | Cell Signaling Technologies | 2217 | 1:1000 |
| Anti-pS235/236-S6 | Cell Signaling Technologies | 2211 | 1:1000 |
| Anti-V5-tag | Cell Signaling Technologies | 13202 | 1:5000 |
| Anti-Vinculin | Sigma-Aldrich | V9131 | 1:10000 |

#### Secondary antibodies

| **Antibody** | **Supplier** | **Catalog #** | **Dilution Used** |
| --- | --- | --- | --- |
| Goat anti-Rabbit IgG – IRDye 800CW | Li-Cor Biosciences | 926-32211 | 1:10000 |
| Goat anti-Mouse IgG – IRDye 680LT | Li-Cor Biosciences | 926-68020 | 1:10000 |

### CMML colony forming assays and xenotransplantation experiments

#### Patient samples

Mononuclear cells were obtained from bone marrow aspirates from 4 distinct CMML patients seen at the Moffitt Cancer Center (Tampa, FL). Patient samples were collected after obtaining written informed consent. Somatic mutations were identified using a targeted gene sequencing panel at the Moffitt Cancer Center (60). The use of human materials was approved by the Institutional Review Board of the Moffitt Cancer Center Scientific Review Committee and the University of South Florida Institutional Review Board in accordance with the Declaration of Helsinki. Additional patient and disease characteristics are provided in Supplementary Tables 7 and 8.

#### Colony forming assays

Viably frozen bone marrow mononuclear cells were the source material for *in vitro* colony forming assays, which were performed as previously described (34). Briefly, bone marrow mononuclear cells were washed in Iscove’s modified Dulbecco’s medium (Lonza, 12-722F) containing 10% fetal bovine serum and subsequently resuspended at a concentration of 5×10^5^ cells/mL in 4-5mL of MethoCult H4230 methylcellulose (STEMCELL Technologies, 04230) containing penicillin/streptomycin, 10ng/mL GM-CSF (R&D Systems, 215-GM) with 250nM dasatinib (Selleck Chemicals, S7782) or DMSO. Next, 1.1mL of the MethoCult cell suspension was plated in 12-well plates in triplicate or quadruplicate. Plates were placed into an incubator at 37°C/5% CO2 for 14 days, and colony counts were determined using a STEMvision automated analyzer (STEMCELL Technologies, Cambridge, MA).

#### Xenotransplantation

All animal experiments were carried out in accordance with institutional guidelines approved by the Moffitt Cancer Center. NOD.Cg-*Prkdc^scid^ Il2rg^tm1Wjl^* Tg(CMV-IL3,CSF2,KITLG)1Eav/MloySzJ (NSG-S) mice were purchased from the Jackson Laboratory (stock no. 013062) and bred under pathogen-free conditions. Viably frozen bone marrow mononuclear cells were the source material for xenografts, which were generated as previously described (34). Additional details regarding the number of cells injected per xenograft model, number of mice per group, pre-injection T cell depletion of the graft, and timing of dasatinib treatment initiation are provided in Supplementary Table 9. Briefly, NSG-S mice (6-10 weeks old) were sublethally irradiated (250 cGy, Mark I Gamma Irradiator, J.L. Shephard, San Fernando, CA) 1 day prior to tail vein injection of ∼3-7×10^6^ bone marrow mononuclear cells. After 2-4 weeks of engraftment, mice were treated by oral gavage with 50mg/kg dasatinib or vehicle (4% DMSO, 30% PEG 300, 5% Tween80 in sterile distilled water) over a 2-week period on weekdays. Spleen and bone marrow were harvested after completion of the treatment course. Spleens were weighed. Enumeration of human CD45^+^ cells in spleen and bone marrow was performed by flow cytometry after staining with BV605-conjugated human-specific CD45 antibody (clone HI30, BD Biosciences, San Jose, CA) using a BD LSRII flow cytometer (BD Biosciences, San Jose, CA). Flow cytometry data were analyzed using FlowJo software version 10 (FlowJo, Ashland, OR). Human CD45 immunohistochemistry was performed by the Brigham and Women’s Hospital Pathology Core using anti-CD45, clone M0701 (Agilent Dako, Santa Clara, CA). Images were captured using a Nikon Eclipse 90i microscope.

### Graphs, Tables, and Figures

All graphs were generated using GraphPad Prism version 7 (San Diego, CA). Morpheus software (https://software.broadinstitute.org/morpheus) was used to perform the nearest neighbor analysis and generate the heatmap and statistics in Figure 3A. Tables were generated in Microsoft Excel or PowerPoint (Redmond, WA). Figures were prepared using Microsoft PowerPoint.

## Acknowledgments

We thank members of the Ebert lab for helpful discussions. This work was supported by the NIH (T32HL066987) and the American Society of Hematology – Amos Medical Faculty Development Program Award to R.B.; NIH (T32GM007753 and F30CA236112 to S.K.); NIH (U24CA210986, U01CA214125, and U24DK112340) to S.A.C.; NIH (R37CA234021) to E.P.; NIH (R01HL082945, P01CA108631, and P50CA206963), the Howard Hughes Medical Institute, the Edward P. Evans Foundation, the Leukemia and Lymphoma Society, and the Janna Brown Charitable Trust to B.L.E. B.L.E. has received research funding from Celgene and Deerfield and serves on the scientific advisory boards for Skyhawk Therapeutics and Exo Therapeutics.

## Author Contributions

R.B. and B.L.E. conceived the study. R.B., S.H.J.K., N.D.U., A.V., M.S., E.P. and B.L.E. designed the experiments. R.B., S.H.J.K., N.D.U., A.V., L.S., T.S., C.H., C.S., and M.S. performed the experiments and/or analyzed the data. N.D.U., M.S., S.A.C., E.P. and B.L.E. supervised the work. R.B., S.H.J.K., and B.L.E. drafted the manuscript. N.D.U., S.A.C., and E.P. edited and provided feedback on the manuscript. All authors reviewed the final version of the manuscript.

## Supplementary Tables

**Supplementary Table 1. Experimental setup for relative quantitation of the total proteome and phosphoproteome in 32D cell lines expressing WT or mutant CBL**

**Supplementary Table 2. Relative quantitation of the total proteome in 32D cell lines expressing WT or mutant CBL**

**Supplementary Table 3. Relative quantitation of proteome-normalized, tyrosine-phosphorylated peptides in 32D cell lines expressing WT or mutant CBL**

**Supplementary Table 4. Relative quantitation of proteome-normalized, serine- and threonine-phosphorylated peptides in 32D cell lines expressing WT or mutant CBL**

**Supplementary Table 5. Experimental setup for relative quantitation of CBL-interacting proteins in 32D cell lines expressing WT or mutant CBL**

**Supplementary Table 6. Relative quantitation of CBL-interacting proteins in 32D cell lines expressing WT or mutant CBL**

**Supplementary Table 7. Mutations and variant allele fractions detected by targeted sequencing of human CMML samples**

**Supplementary Table 8. Clinical, pathologic, and laboratory characteristics of CMML patients at the time of bone marrow sample collection**

**Supplementary Table 9. CMML xenotransplantation experimental details**

## Supplementary Figure Legends

**Supplementary Figure 1. Characteristics of *CBL* mutations in 191 patients**

(A) Position and type of *CBL* mutations detected within exons 7-9, which include CBL’s linker region and RING domain. (B) Distribution of *CBL* mutations by variant type. (C) Number of missense mutations detected at each amino acid position encoded within *CBL* exons 7-9.

**Supplementary Figure 2. Generation of Cbl knockout cell lines by CRISPR-Cas9-mediated gene editing in 32D cells.**

(A) *Cbl* exon 1 targeting by CRISPR-Cas9. (B) Western blot for CBL in single cell clones expressing Cas9 and gRNA targeting *Cbl* exon 1. Polyclonal antibody used for detection of CBL (Santa Cruz Biotechnology, sc170) by western blot recognizes an epitope on the C-terminus of the protein. *Cbl* knockout clone 5A3 was used for experiments.

**Supplementary Figure 3. Expression of CBL mutants confers IL3-independence and a competitive proliferative advantage in BaF3 cells.**

(A) Viable cell counts after IL3 withdrawal from BaF3-*Cbl*^KO^ cells expressing CBL WT, Y371H, C384Y, or R420Q. (B) Competition between GFP-labeled cells expressing CBL WT or C384Y and dTomato-labeled cells expressing CBL WT. Cells were initially mixed at ratio of 1:10 GFP:dTomato and assessed by flow cytometry every 3-6 days.

**Supplementary Figure 4. Targeting splice sites in *Cbl* introns 7 and 8 leads to exon 8 exclusion.**

(A) *Cbl* intron gRNAs targeting exon 8 splice sites. (B) PCR amplification and sequencing of *Cbl* exon 8 in 32D cells expressing Cas9 and a non-targeting gRNA (NT) or gRNAs targeting *Cbl* exon 8 splice sites.

**Supplementary Figure 5. Expression of CBL C384Y is associated with increased CBL phosphorylation, LYN activation and PI3K/AKT signaling in TF1 cells.**

Western blot for total and phosphorylated CBL, LYN, AKT and S6 protein in TF1-*CBL*^KO^ cells expressing V5-tagged CBL WT or C384Y. Western blot for vinculin (VCL) was used as a loading control.

**Supplementary Figure 6. Lyn binding to the proline-rich region of CBL C384Y is increased in TF1 cells.**

Western blot for V5, LYN and vinculin (VCL) in anti-V5 IP samples and whole-cell lysates (WCL) from TF1-*CBL*^KO^ cells expressing V5-tagged CBL WT, C384Y, or C384Y/**Δ**PRR.

**Supplementary Figure 7. 32D*-Lyn*^KO^ cells are at a proliferative disadvantage in competition against 32D-*Lyn*^WT^ cells.**

Competition between CBL C384Y-expressing 32D cells on *Lyn*^WT^ (dTomato) and *Lyn*^KO^ (GFP) genetic backgrounds.

**Supplementary Figure 8. 32D*-Lyn*^KO^ cells expressing CBL C384Y show reduced phosphorylation of CBL and AKT**

(A) Western blot for total and phosphorylated CBL, LYN, AKT, and S6 proteins in 32D-*Cbl*^KO^ cells expressing V5-tagged CBL WT, Y371H, C384Y, or R420Q. Western blot for **β**-actin (ACTB) was used as a loading control. (B) Quantitation of pS473-AKT/AKT by densitometry from 4 independent experiments. Bars depict mean and SD.

**Supplementary Figure 9. Dasatinib inhibits proliferation of *CBL* mutant TF1 cells and 32D cells with CRISPR-Cas9 targeted mis-splicing of *Cbl* exon 8**

(A) Proliferation of TF1-*CBL*^KO^ cells expressing luciferase (KO) or V5-tagged CBL WT or C384Y in the presence of DMSO or a range of dasatinib concentrations over 3 days. (B and C) Competition between dTomato-labeled TF1-*CBL*^KO^ cells expressing CBL WT and GFP-labeled TF1-*CBL*^KO^ cells expressing (B) CBL C384Y or (C) CBL R420Q in the presence of DMSO (closed symbols) or 1μM dasatinib (open symbols). (D and E) Competition between 32D-Cas9 cells expressing *Cbl* intron (D) 7-1 or (E) 7-2 gRNAs (RFP657) and 32D-Cas9 cells expressing non-targeting gRNA (BFP) in the presence of DMSO (closed symbols) or 1μM dasatinib (open symbols).

**Supplementary Figure 10. Dasatinib inhibits CBL phosphorylation, LYN activation and downstream PI3K/AKT signaling in TF1 cells expressing CBL C384Y**

Western blot for total and phosphorylated CBL, LYN, AKT and S6 proteins in TF1-*CBL*^KO^ cells expressing V5-tagged CBL WT or C384Y treated with 1μM dasatinib for 2 hours. Western blot for vinculin (VCL) was used as a loading control.

**Supplementary Figure 11. Effect of dasatinib treatment on the competitive advantage of CBL C384Y-expressing 32D cells on *Lyn*^WT^ and *Lyn*^KO^ genetic backgrounds**

(A-C) Competition between CBL WT (dTomato) and C384Y (GFP) expressing 32D cells on *Lyn*^WT^ (A) and *Lyn*^KO^ (B and C) genetic backgrounds in the presence of DMSO (closed symbols) or 1μM dasatinib (open symbols).

**Supplementary Figure 12. Effect of dasatinib treatment on spleen weight and bone marrow expansion of CMML cells in NSG-S mice**

(A) Spleen weights in vehicle- and dasatinib-treated mice. (B) Percent change in spleen weights in vehicle- and dasatinib mice. (C) Percentage of human CD45^+^ cells in the bone marrow of vehicle- and dasatinib-treated mice (D) Percent change in human CD45^+^ cells in vehicle- and dasatinib-treated mice. Each dot represents an individual mouse. Bars depict mean and S.D.

## Notes

**Financial Support:** NIH (T32HL066987) and the American Society of Hematology – Amos Medical Faculty Development Program Award to R.B.; NIH (T32GM007753 and F30CA236112) to S.H.J.K.; NIH (U24CA210986, U01CA214125, and U24DK112340) to S.A.C.; NIH (R37CA234021) to E.P.; NIH (R01HL082945, P01CA108631, and P50CA206963), the Howard Hughes Medical Institute, the Edward P. Evans Foundation, the Leukemia and Lymphoma Society, and the Janna Brown Charitable Trust to B.L.E.

**Conflict of Interest Disclosure:** B.L.E. has received research funding from Celgene and Deerfield and serves on the scientific advisory boards for Skyhawk Therapeutics and Exo Therapeutics. The remaining authors declare no potential conflicts of interest.

