## Supplementary Data for "CBL mutations promote activation of PI3K/AKT signaling via LYN kinase"

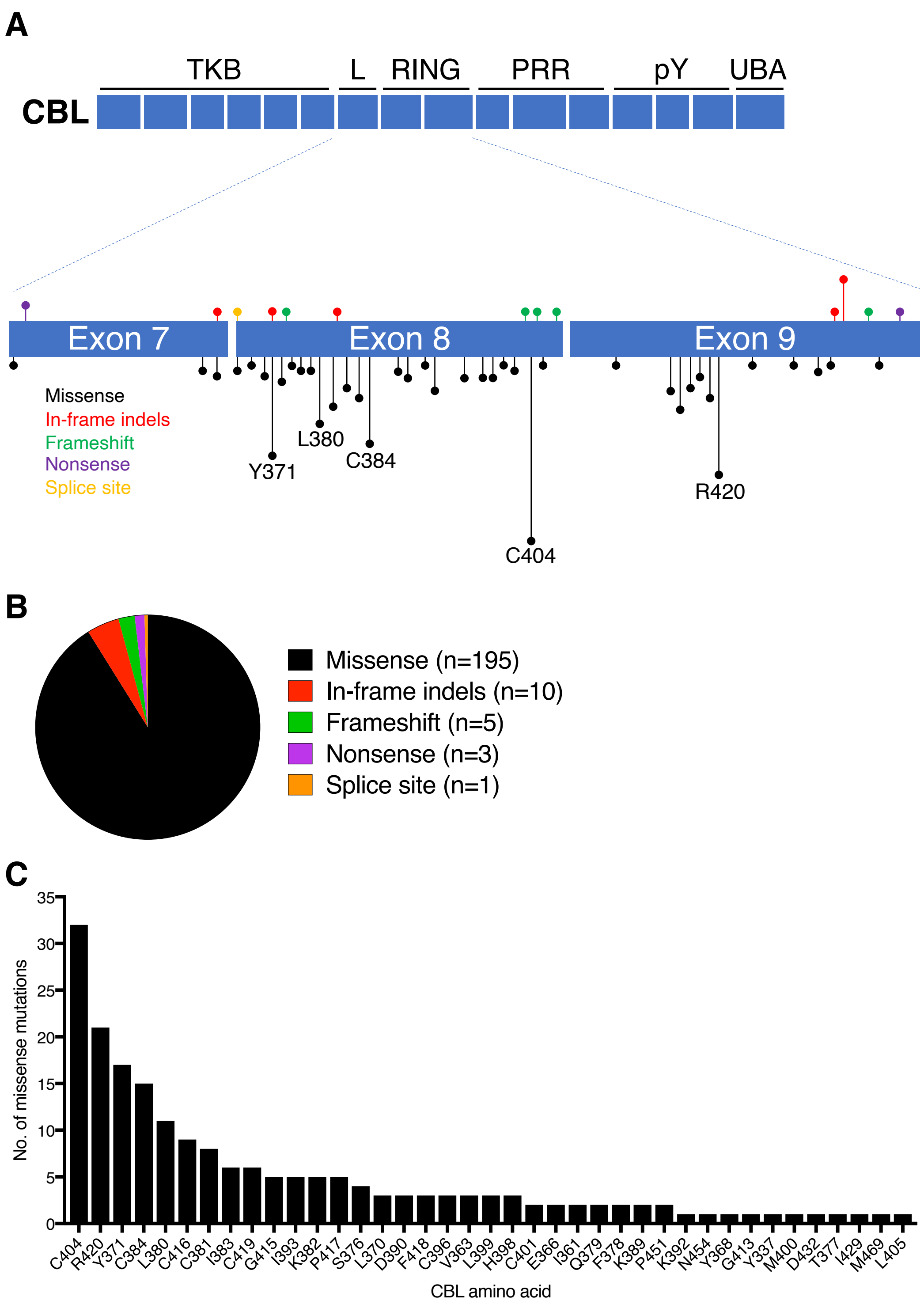


**Supplementary Figure 1. Characteristics of *CBL* mutations in 191 patients.**
(A) Position and type of *CBL* mutations detected within exons 7-9, which includes CBL's linker region (L) and RING domain. (B) Distribution of *CBL* mutations by variant type. (C) Number of missense mutations detected at each amino acid position encoded with *CBL* exons 7-9.

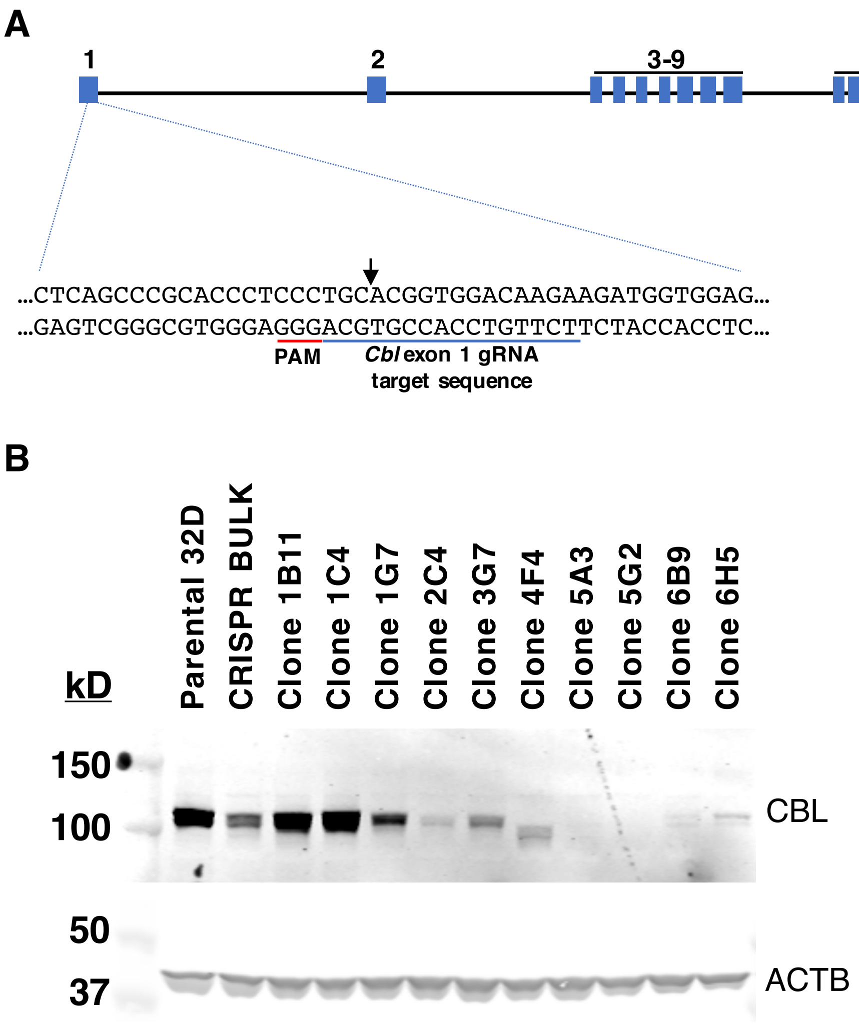


**Supplementary Figure 2. Generation of *Cbl* knockout cell lines by CRISPR-Cas9 mediated gene editing in 32D cells.**
(A) *Cbl* exon 1 targeting by CRISPR-Cas9. (B) Western blot for CBL in single cell clones expressing Cas9 and gRNA targeting *Cbl* exon 1. Polyclonal antibody used for detection of CBL by western blot recognizes an epitope on the C-terminus of the protein (Santa Cruz Biotechnology, sc170). Western blot for **β**-actin (ACTB) was used as a loading control. *Cbl* knockout clone 5A3 was used for experiments.

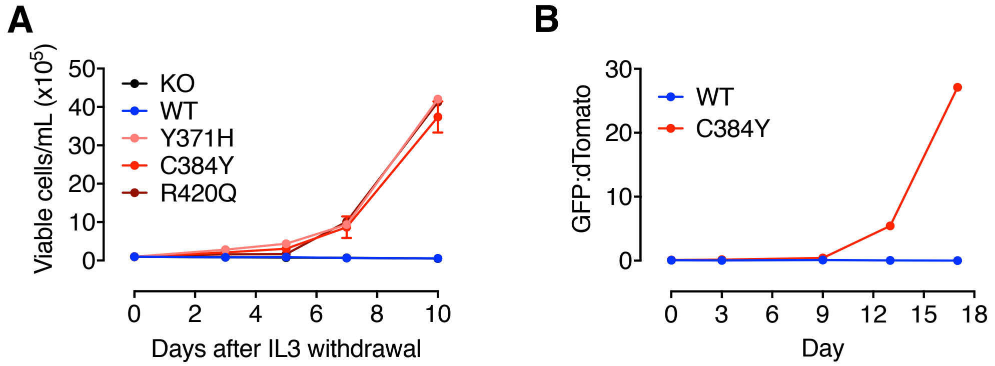


**Supplementary Figure 3. Expression of CBL mutants confers IL3-independence and a competitive proliferative advantage in BaF3 cells.**
(A) Viable cell counts after IL3 withdrawal from BaF3-*Cbl*$^{KO}$ cells expressing CBL WT, Y371H, C384Y, or R420Q. Mean and S.D. of triplicate values are depicted. (B) Competition between GFP-labeled cells expressing CBL WT or C384Y and dTomato-labeled cells expressing CBL WT. Cells were initially mixed at ratio of 1:10 GFP:dTomato and assessed by flow cytometry every 3-6 days. Symbols depict mean of duplicate values.

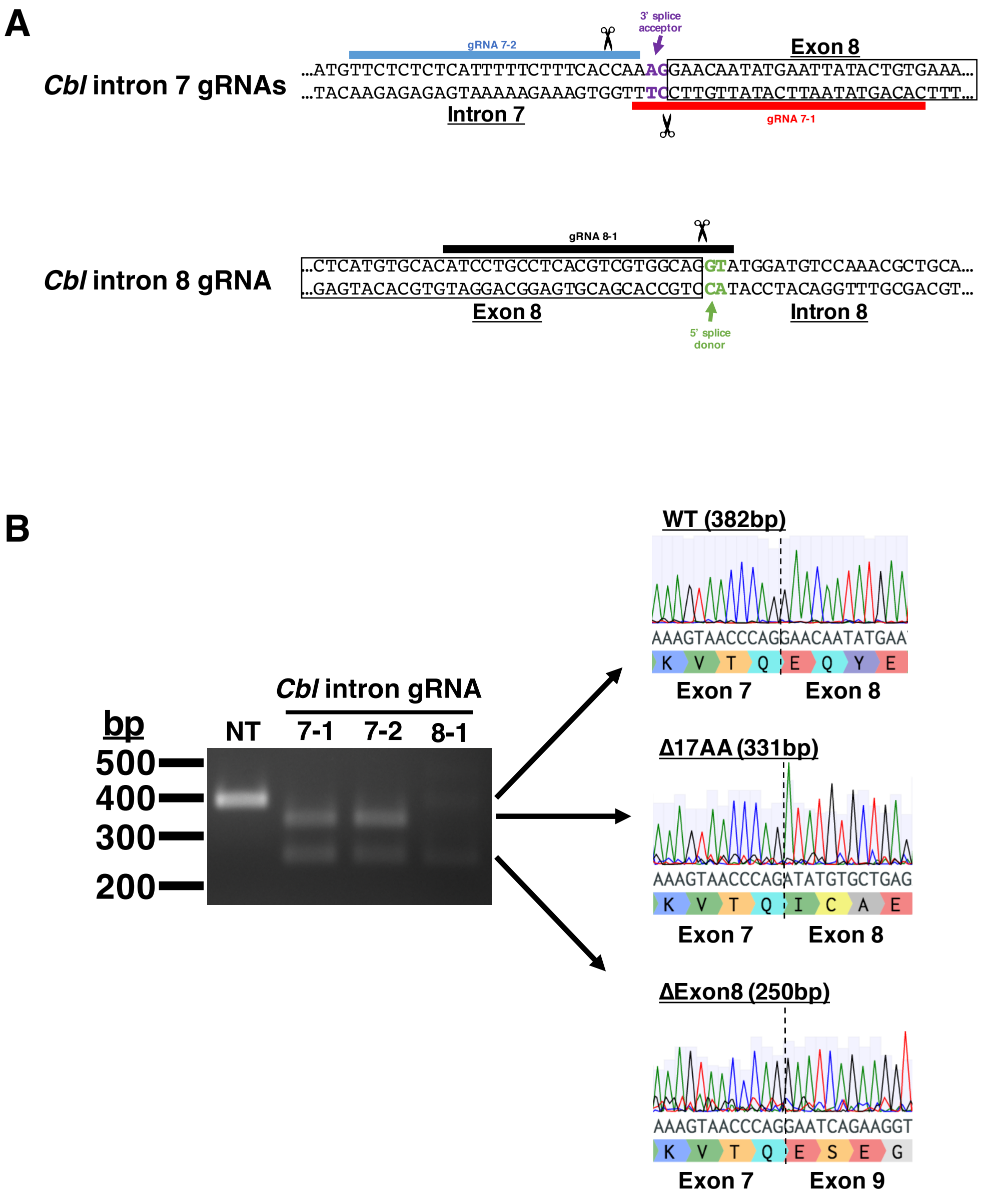


**Supplementary Figure 4. Targeting splice sites in Cbl introns 7 and 8 leads to exon 8 exclusion.** (A) *Cbl* intron gRNAs targeting exon 8 splice sites. (B) PCR amplification and sequencing of *Cbl* exon 8 in 32D cells expressing Cas9 and a non-targeting gRNA (NT) or gRNAs targeting *Cbl* exon 8 splice sites.

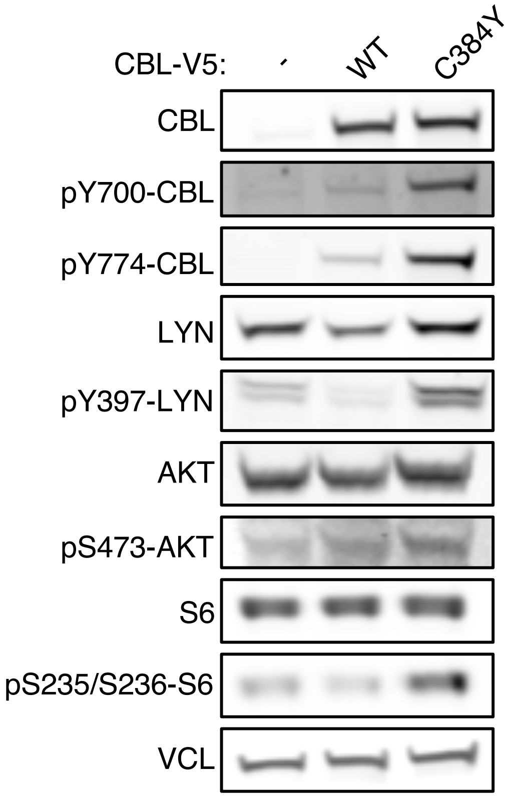


**Supplementary Figure 5. Expression of CBL C384Y is associated with increased CBL phosphorylation, LYN activation, and PI3K/AKT signaling in TF1 cells.**
Western blot for total and phosphorylated CBL, LYN, AKT, and S6 protein in TF1-*CBL*$^{KO}$ cells expressing V5-tagged CBL WT or C384Y. Western blot for vinculin (VCL) was used as a loading control.

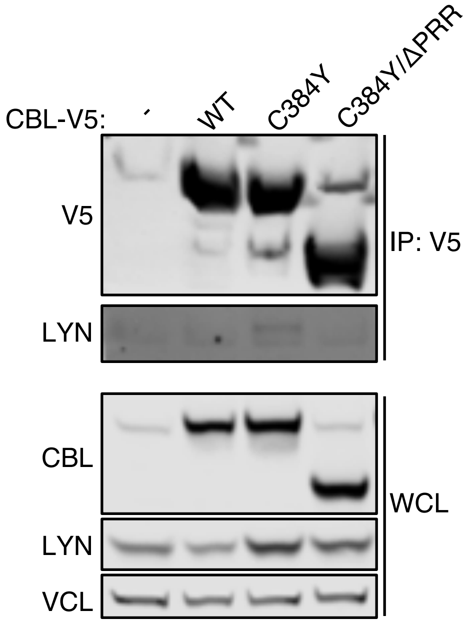


**Supplementary Figure 6. Lyn binding to the proline-rich region of CBL C384Y is increased in TF1 cells.**
Western blot for V5, LYN and vinculin (VCL) in anti-V5 IP samples and whole-cell lysates (WCL) from TF1 cells expressing V5-tagged CBL WT, C384Y, or C384Y/ΔPRR.

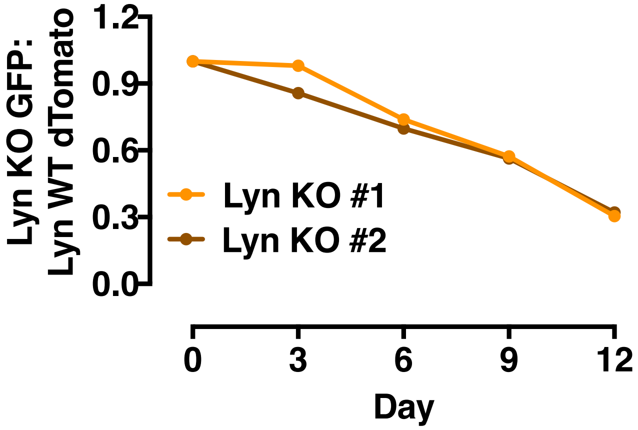


**Supplementary Figure 7. 32D-*Lyn*$^{KO}$ cells are at a proliferative disadvantage in competition against 32D-*Lyn*$^{WT}$ cells.**
Competition between CBL C384Y-expressing 32D cells on *Lyn*$^{WT}$ (dTomato) and *Lyn*$^{KO}$ (GFP) genetic backgrounds. Symbols depict the mean of duplicates values.

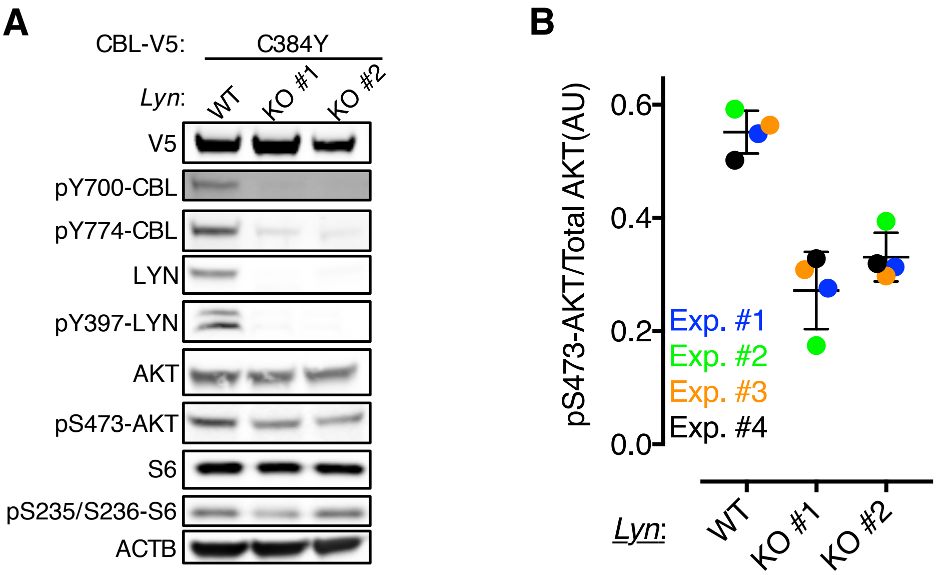


**Supplementary Figure 8. 32D-*Lyn*$^{KO}$ cells expressing CBL C384Y show reduced phosphorylation of CBL and AKT.**

(A) Western blot for total and phosphorylated CBL, LYN, AKT, and S6 proteins in 32D-*Cbl*$^{KO}$ cells expressing V5-tagged CBL WT, Y371H, C384Y, or R420Q. Western blot for **β**-actin (ACTB) was used as a loading control. (B) Quantitation of pS473-AKT/AKT by densitometry from 4 independent experiments. Bars depict mean and S.D.

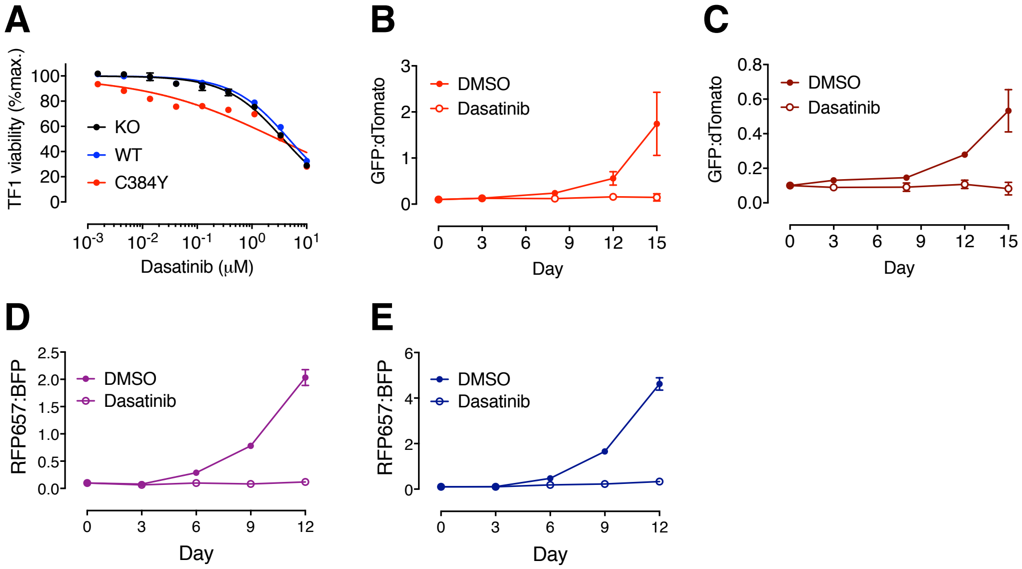


**Supplementary Figure 9. Dasatinib inhibits proliferation of *CBL* mutant TF1 cells and 32D cells with CRISPR-Cas9 targeted mis-splicing of *Cbl* exon 8.**

(A) Proliferation of TF1-*CBL*$^{KO}$ cells expressing luciferase (KO) or V5-tagged CBL WT or C384Y in the presence of DMSO or a range of dasatinib concentrations over 3 days. (B and C) Competition between dTomato-labeled TF1-*CBL*$^{KO}$ cells expressing CBL WT and GFP-labeled TF1-*CBL*$^{KO}$ cells expressing (B) CBL C384Y or (C) CBL R420Q in the presence of DMSO (closed symbols) or 1μM dasatinib (open symbols). (D and E) Competition between 32D-Cas9 cells expressing *Cbl* intron (D) 7-1 or (E) 7-2 gRNAs (RFP657) and 32D-Cas9 cells expressing non-targeting gRNA (BFP) in the presence of DMSO (closed symbols) or 1μM dasatinib (open symbols). Mean and S.D. of triplicate values are depicted.

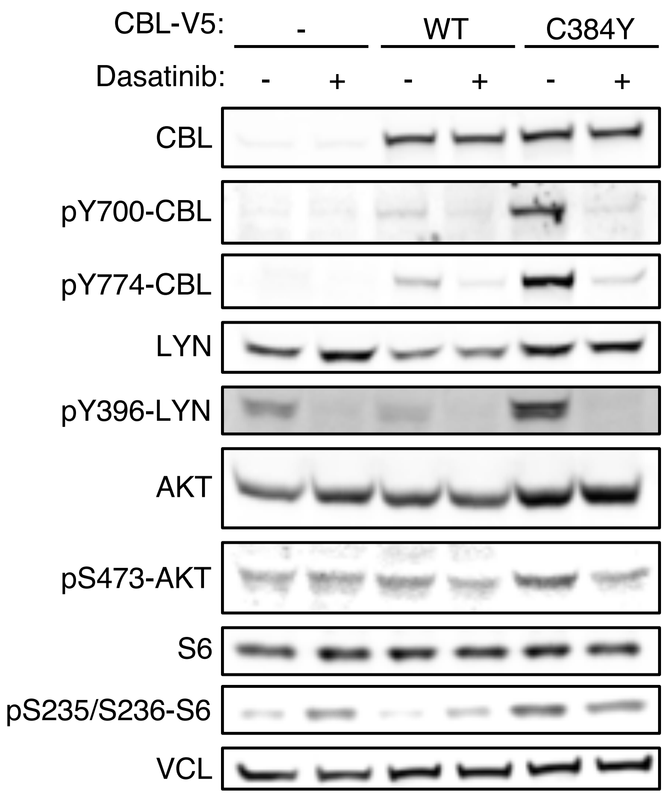


**Supplementary Figure 10. Dasatinib inhibits CBL phosphorylation, LYN activation and downstream PI3K/AKT signaling in TF1 cells expressing CBL C384Y.**
Western blot for total and phosphorylated CBL, LYN, AKT, and S6 proteins in TF1-*CBL*$^{KO}$ cells expressing V5-tagged CBL WT or C384Y after treatment with 1μM dasatinib or DMSO for 2 hours. Western blot for vinculin (VCL) was used as a loading control.

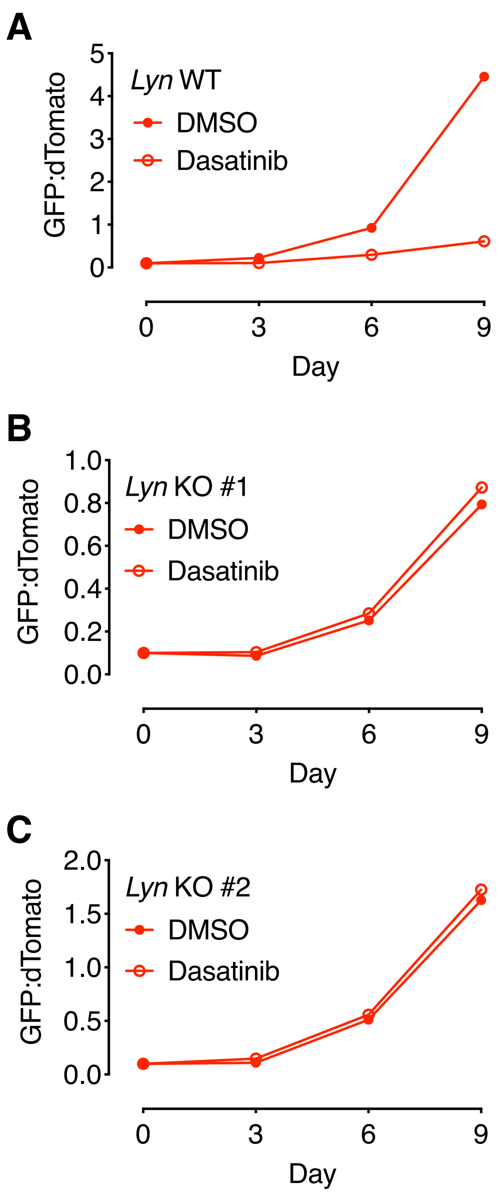


**Supplementary Figure 11. Effect of dasatinib treatment on the competitive advantage of CBL C384Y-expressing 32D cells on *Lyn*$^{WT}$ and *Lyn*$^{KO}$ genetic backgrounds.**
(A-C) Competition between CBL WT (dTomato) and C384Y (GFP) expressing 32D cells on *Lyn*$^{WT}$ (A) and *Lyn*$^{KO}$ (B and C) genetic backgrounds in the presence of DMSO (closed symbols) or 1μM dasatinib (open symbols). Symbols depict the mean of duplicates values.

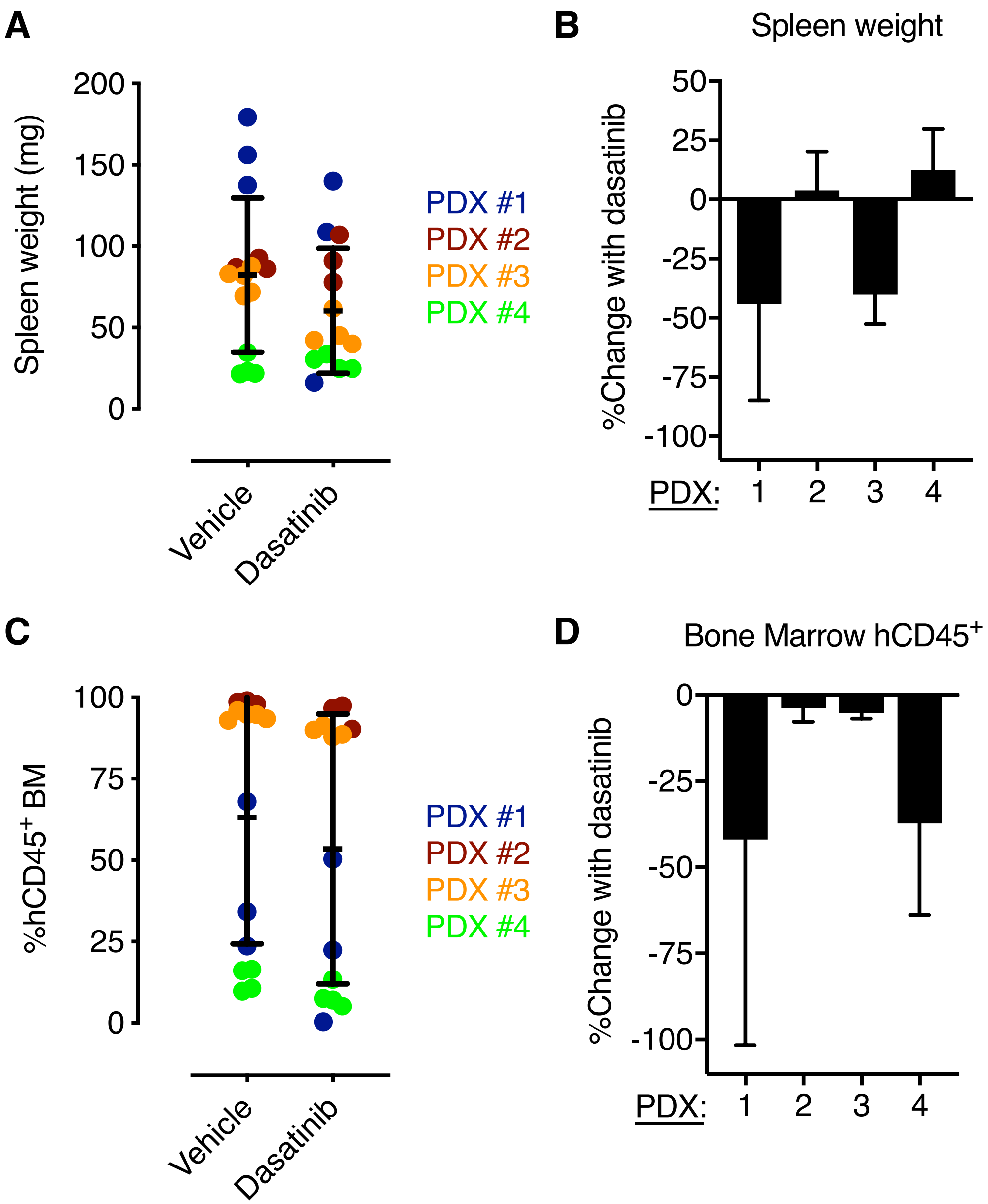


**Supplementary Figure 12. Effect of dasatinib treatment on spleen weight and bone marrow expansion of CMML cells in NSG-S mice.**

(A) Spleen weights in vehicle- and dasatinib-treated mice. (B) Percent change in spleen weights in vehicle- and dasatinib mice. (C) Percentage of human CD45$^{+}$ cells in the bone marrow of vehicle- and dasatinib-treated mice (D) Percent change in human CD45$^{+}$ cells in vehicle- and dasatinib-treated mice. Each dots represents an individual mouse. Bars depict mean and S.D.

**Supplementary Table 7. Mutations and variant allele frequencies detected by targeted sequencing of human CMML samples.**

| CMML PDX | Mutation (variant allele fraction) | | | | | | |
|---|---|---|---|---|---|---|---|
| 1 | CBL R462L (0.16) | CEBPA L139_D140del (0.16) | KDM6A W553L (0.13) | KDM6A D1206Y (0.15) | PHF6 R319L (0.12) | STAG2 L716F (0.25) | STAG2 E721X (0.16) |
| 2 | CBL R420P (0.95) | ASXL1 G646Wfs (0.38) | SETBP1 D868N (0.47) | SRSF2 P95H (0.54) | TET2 C1263Y (0.49) | | |
| 3 | CBL D390V (0.49) | ASXL1 L775X (0.34) | SETBP1 D868N (0.38) | TET2 T556fs (0.33) | TET2 Q562X (0.37) | SRSF2 P95R (0.47) | |
| 4 | CBL H398Q (0.24) | TET2 K244fs (0.51) | SRSF2 P95H (0.57) | SETBP1 I871T (0.76) | | | |

**Supplementary Table 8. Clinical, pathologic, and laboratory characteristics of CMML patients at the time of bone marrow sample collection.**

| CMML PDX | WHO | FAB | Splenomegaly | WBC ($x10^3/\mu L$) | Monocytes ($x10^3/\mu L$) | Hgb (g/dL) | Platelets ($x10^3/\mu L$) | CG |
|---|---|---|---|---|---|---|---|---|
| 1 | CMML-0 | MP | Yes | 32.9 | 3.3 | 10.3 | 311 | 46, XX |
| 2 | CMML-2 | MP | Yes | 46.1 | 5.1 | 10.6 | 146 | 46, XY |
| 3 | CMML-1 | MP | Yes | 128.1 | 1.3 | 8.5 | 49 | 46, XY |
| 4 | CMML-0 | MP | Yes | 42.4 | 1.3 | 12.6 | 211 | 46, XY |

### Supplementary Table 9. CMML xenotransplantation experimental details

| CMML PDX | Mutations | Graft T cell depleted? | Cells/Mouse ($x10^6$) | No. mice treated with dasatinib* | No. mice treated with vehicle* | Treatment initiation (day post-transplant) | Treatment duration# | Endpoint (days post-transplant) |
|---|---|---|---|---|---|---|---|---|
| 1 | CBL, CEBPA, KDM6A, PHF6, STAG2 | No | 4.7 | 3 | 3 | 14 | 2 weeks | 31 |
| 2 | CBL, ASXL1, SETPB1, SRSF2, TET2 | Yes | 2.6 | 3 | 3 | 14 | 2 weeks | 28 |
| 3 | CBL, ASXL1, SETBP1, SRSF2, TET2 | Yes | 6.5 | 4 | 5 | 28 | 2 weeks | 42 |
| 4 | CBL, SETBP1, SRSF2, TET2 | Yes | 3.75 | 4 | 4 | 28 | 2 weeks | 42 |

*Dasatinib was administered by oral gavage at 50mg/kg. Vehicle was composed of 4% DMSO, 30% PEG 300, 5% Tween80 in sterile distilled water.

#Mice were treated daily Monday through Friday over a 2-week period.
